# A White Noise Approach to Evolutionary Ecology

**DOI:** 10.1101/2020.07.28.226001

**Authors:** Bob Week, Scott L. Nuismer, Luke J. Harmon, Stephen M. Krone

## Abstract

Although the evolutionary response to random genetic drift is classically modelled as a sampling process for populations with fixed abundance, the abundances of populations in the wild fluctuate over time. Furthermore, since wild populations exhibit demographic stochasticity, it is reasonable to consider the evolutionary response to demographic stochasticity and its relation to random genetic drift. Here we close this gap in the context of quantitative genetics by deriving the dynamics of the distribution of a quantitative character and the abundance of a biological population from a stochastic partial differential equation driven by space-time white noise. In the process we develop a useful set of heuristics to operationalize the powerful, but abstract theory of white noise and measure-valued stochastic processes. This approach allows us to compute the full implications of demographic stochasticity on phenotypic distributions and abundances of populations. We demonstrate the utility of our approach by deriving a quantitative genetic model of diffuse coevolution mediated by exploitative competition for a continuum of resources. In addition to trait and abundance distributions, this model predicts interaction networks defined by rates of interactions, competition coefficients, or selection gradients. Analyzing the relationship between selection gradients and competition coefficients reveals independence between linear selection gradients and competition coefficients. In contrast, absolute values of linear selection gradients and quadratic selection gradients tend to be positively correlated with competition coefficients. That is, competing species that strongly affect each other’s abundance tend to also impose selection on one another, but the directionality is not predicted. This approach contributes to the development of a synthetic theory of evolutionary ecology by formalizing first principle derivations of stochastic models that underlie rigorous investigations of the relationship between feedbacks of biological processes and the patterns of diversity they produce.

## 1. Introduction

Current mathematical approaches to synthesize the dynamics of abundance and evolution in populations have capitalized on the fact that biological fitness plays a key role in determining both sets of dynamics. In particular, while covariance of fitness and genotype is the basis of evolution by natural selection, the mean fitness across all individuals in a population determines the growth, stasis or decline of abundance. Although this connection has been established in the contexts of population genetics (Crow and Kimura, 1970, Roughgarden, 1979), evolutionary game theory (Hofbauer and Sigmund, 1998, Lion, 2018, Nowak, 2006), quantitative genetics (Doebeli, 1996, Lande, 1982, Lion, 2018) and a unifying framework for these three distinct approaches to evolutionary theory (Champagnat et al., 2006), there remains a gap in incorporating the intrinsically random nature of abundance into the evolution of continuous traits. Specifically, in theoretical quantitative genetics the derivation of a population’s response to random genetic drift is derived in discrete time under the assumption of constant effective population size using arguments based on properties of random samples (Lande, 1976). Though this approach conveniently mimics the formalism provided by the Wright-Fisher model of population genetics, real population sizes fluctuate over time. Furthermore, since these fluctuations are themselves stochastic, it seems natural to derive expressions for the evolutionary response to demographic stochasticity and consider how the results relate to characterizations of random genetic drift. This can be done in continuous time for population genetic models without too much technical overhead, assuming a finite number of alleles (Gomulkiewicz et al., 2017, Lande et al., 2009, Parsons et al., 2010). However, for populations with a continuum of types, such as a quantitative trait, finding a formal approach to derive the evolutionary response to demographic stochasticity has remained a vexing mathematical challenge. In this paper we close this gap by combining the calculus of white noise with results on rescaled limits of measure-valued branching processes (MVBP) and stochastic partial differential equations (SPDE).

Our goals in this paper are twofold: 1) Establish a novel synthetic approach to theoretical evolutionary ecology that provides a formal connection between demographic stochasticity and random genetic drift in the context of quantitative traits. 2) Communicate some useful properties of space-time white noise, MVBP and SPDE to a wide audience of mathematical evolutionary ecologists. With these goals in mind we will not provide a rigorous treatment of any of these mathematically rich topics. Instead, we introduce a set of heuristics that only require the basic concepts of Riemann integration, partial differentiation and some exposure to Brownian motion and stochastic ordinary differential equations (SDE). A concise introduction to SDE and Brownian motion has been provided by Evans (2014).

Since MVBP are abstract mathematical objects and their rigorous study requires elaborate mathematical machinery, the use of MVBP in mainstream theoretical evolutionary ecology has been limited. However, they provide natural models of biological populations by capturing various mechanistic details. In particular, MVBP generalize classical birth-death processes, such as the Galton-Watson process (Kimmel and Axelrod, 2015, Dawson, 1993), to model populations of discrete individuals that carry some value in a given type-space. Selection can then be modelled by associating these values with average reproductive output and mutation can be incorporated using a model that determines the distribution of offspring values given their parental value. For population genetic models the type-space is the discrete set of possible alleles individuals can carry. In quantitative genetic models tracking the evolution of *d*-dimensional phenotypes, this type-space is typically set to the Euclidean space 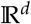. By starting with branching processes we can implement mechanistic models of biological fitness that account for the phenotype of the focal individual along with the phenotypes and number of all other individuals in a population or community. By taking a rescaled limit, we can then use these detailed individual-based models to derive population-level models tracking the dynamics of population abundance and phenotypic distribution driven by selection, mutation and demographic stochasticity. Hence, rescaled limits of MVBP provide a means to derive mathematically tractable, yet biologically mechanistic models of eco-evolutionary dynamics.

For univariate traits (i.e., *d* = 1) Konno and Shiga (1988), Reimers (1989), Li (1998) and Champagnat et al. (2006) have shown that rescaled limits for a large class of MVBP converge to solutions of SPDE. Although cases in which *d* ≥ 1 can be treated using the so-called martingale problem formulation (Dawson, 1993), the SPDE formulation provides a more intuitive description of the biological processes involved. We therefore focus on the case *d* = 1 here. This allows us to introduce a concrete set of heuristics for deriving SDE tracking the dynamics of abundance, phenotypic mean and phenotypic variance to a wide audience of mathematical evolutionary ecologists. Following our approach to simplify notation and develop heuristics for calculations, future work can possibly use the martingale formulation to extend the results presented here for *d* > 1 and even for infinite-dimensional traits (Dawson, 1993, Stinchcombe et al., 2012). Rigorous introductions to SPDE and rescaled limits of MVBP have been respectively provided by Da Prato and Zabczyk (2014) and Etheridge (2000).

In this paper we begin in §2 by introducing the basic framework of our approach. We first outline the essential ideas behind deriving evolutionary dynamics from abundance dynamics using a deterministic partial differential equation (PDE). In SM §3.1 we review rescaled limits of MVBP, their associated SPDE and introduce an approach to derive SDE tracking the dynamics of abundance, phenotypic mean and phenotypic variance. This approach requires performing calculations with respect to space-time white noise processes and we provide heuristics for doing so in SM §2.1. In §2.2 we discuss consequences of the derived SDE for general phenotypic distributions and simplify their expressions by assuming normally distributed phenotypes. For added biological relevance, we incorporate models of inheritance and development following classical quantitative genetics. To demonstrate how our framework can be used to formulate a synthetic theory of evolutionary ecology, in §3 we derive a model of diffuse coevolution for a set of *S* species competing along a resource continuum. The basic approach follows classical niche theory to develop biological fitness as a function of niche parameters and niche locations of other individuals in the community. We then use this model to derive formula for selection gradients and competition coefficients. Finally, we investigate the relationship between selection gradients and competition coefficients using a high-richness (large *S*) approximation.

## 2. The Framework

At the core of our approach is a model of stochastic abundance dynamics for a structured population in continuous time and phenotypic space. From this stochastic equation we derive a system of SDE for the dynamics of total abundance, mean trait and additive genetic variance of a population. In particular, our approach develops a quantitative genetic theory of evolutionary ecology. A popular alternative to quantitative genetics is the theory of adaptive dynamics (Dieckmann and Law, 1996, Metz et al., 1996). As demonstrated by Page and Nowak (2002) and Champagnat et al. (2006), the canonical equation of adaptive dynamics can be derived from the replicator-mutator equation, which in turn can be derived from models of abundance dynamics, revealing a synthesis of mathematical approaches to theoretical evolutionary ecology. In this section we briefly outline derivations of the replicator-mutator equation and trait dynamics from abundance dynamics in the deterministic case. We then extend these formula along with related results to the case of random reproductive output (i.e., demographic stochasticity).

### 2.1. Deterministic Dynamics

#### Finite Number of Types

We start by considering the dynamics of an asexually reproducing population in a homogeneous environment. For simplicity, we first assume individuals are haploid and carry one of *K* alleles each with a different fitness expressed as growth rate before introducing a model involving a quantitative trait. Under these assumptions, the derivation of the evolution of allele frequencies due to natural selection can be derived from expressions of exponential growth. This, and a few related approaches, have been provided by Crow and Kimura (1970). Mutation can be included using a matrix of transition rates. Specifically, denoting *ν_i_* the abundance of individuals with allele *i, m_i_* the growth rate of allele *i* (called the Malthusian parameter in Crow and Kimura, 1970), *μ_ij_* the mutation rate from allele *i* to allele *j* and assuming selection and mutation are decoupled (Bürger, 2000), we have

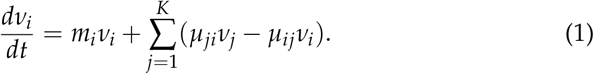

Starting from this model, we get the total abundance of the population as 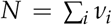, the frequency of allele *i* as *p_i_* = *v_i_/N* and the mean Malthusian fitness of the population as 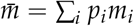. Note we have used the abbreviation 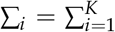 to simplify inline notation. Observing 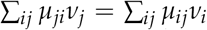, we use linearity of differentiation to derive the dynamics of abundance *dN/dt* as

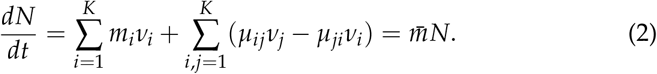

To derive the dynamics of the allele frequencies *p*_1_,…, *p_K_*, we use the quotient rule of elementary calculus to find

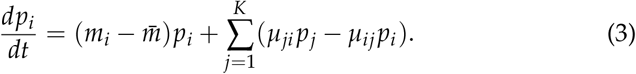

Two important observations of these equations include: (i) Mean Malthusian fitness 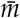 is equivalent to the population growth rate and thus determines the abundance dynamics of the entire population. (ii) Selection for allele *i* occurs when 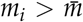 and selection against allele *i* occurs when 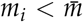. Hence, as mentioned in the introduction, fitness plays a key role in determining both abundance dynamics and evolution.

Equation (3) is known in the field of evolutionary game theory as a replicator-mutator equation (Nowak, 2006). Instead of being explicitly focused on alleles, the replicator-mutator equation describes the fluctuations of relative abundances of various <u>types</u> in a population in terms of replication and annihilation rates of each type and hence can be used to model dynamical systems outside of evolutionary biology (Nowak, 2006).

#### Continuum of Types

Inspired by equations (1)–(3), we derive an analog of the replicator-mutator equation for a continuum of types (that is, for a quantitative trait). In particular, we model a continuously reproducing population with trait values 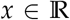 and an abundance density *v*(*x, t*) that represents the amount of individuals in the population with trait value *x* at time t. Hence, the abundance density satisfies 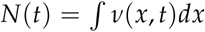 and *p*(*x, t*) = *v*(*x, t*)/*N*(*t*) is the relative density of trait *x* which we also refer to as the phenotypic distribution. Note we have used the abbreviation 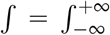 to simplify inline notation.

In analogy with the growth rates *m_i_* for equation (1) we write *m*(*v, x*) as the growth rate associated with trait value *x* which depends on the abundance density *v*. We assume mutation is captured by diffusion with coefficient 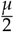. Hence, we model the demographic dynamics of a population and the dynamics of a quantitative character simultaneously by the PDE

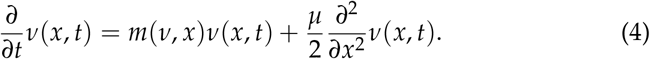

Equation (4) qualifies both as a semilinear evolution equation and also a scalar reaction-diffusion equation. Although the general theory of such equations is quite rich, it is also quite difficult (Evans, 2010, Zheng, 2004). Hence, to stay within the realms of analytical tractability and biological plausibility, we require a set of technical assumptions which we list in SM §1.1. These assumptions guarantee solutions to equation (4) exist for all finite time *t* > 0 and, hence, let us investigate the ecological and evolutionary dynamics of biological populations.

Equation (4) can be seen as an analog of equation (1) for a continuum of types. By assuming mutation acts via diffusion, the effect of mutation causes the abundance density *v*(*x, t*) to flatten out across phenotypic space. In fact, if the growth rate is constant across *x*, then this model of mutation will cause *v*(*x, t*) to converge to a flat line in *x* as *t* → ∞. Interpreting the trait value *x* as location in geographic space, equation (4) becomes a well-studied model of spatially distributed population dynamics (Cantrell and Cosner, 2004).

Although clearly an idealized representation of biological reality, this model is sufficiently general to capture a large class of dynamics including density dependent growth and frequency dependent selection. As an example, logistic growth combined with stabilizing selection can be captured using the growth rate

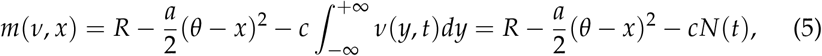

where *a* > 0 the is strength of abiotic stabilizing selection around the phenotypic optimum *θ, c* > 0 is the strength of intraspecific competition and we refer to *R* as the innate growth rate (see §3.3 below). In the language of population ecology, 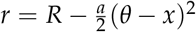 is the intrinsic growth rate of the population (Chesson, 2000). This model assumes competitive interactions cause the same reduction in fitness regardless of trait value.

This exemplary fitness function has a few convenient properties. First, the effect of competition induces a local carrying capacity on the population, leading to a finite equilibrium abundance over bounded subsets of phenotypic (or geographic) space. Second, abiotic selection prevents the abundance density from diffusing too far from the abiotic optimum. In particular, when 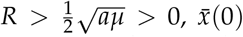 is finite, *σ*^2^(0) is non-negative and finite and *N*(0) is positive and finite, this leads to a unique stable equilibrium given by

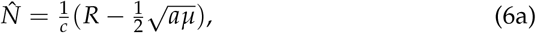

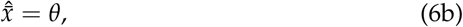

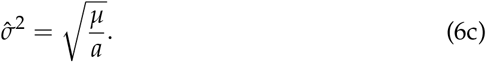

We demonstrate this result in SM §1.2. The equilibrial phenotypic variance predicted by this model coincides with a classic quantitative genetic result predicted by modelling the combined effects of Gaussian stabilizing selection and the Gaussian allelic model of mutation (Bürger, 2000, Johnson and Barton, 2005, Lande, 1975, Walsh and Lynch, 2018).

To derive a replicator-mutator equation from equation (4), we employ integration-by-parts and the chain rule from calculus. Writing

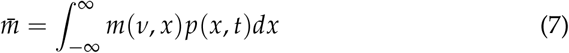

for the mean fitness, we find

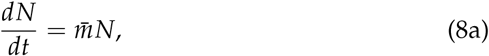

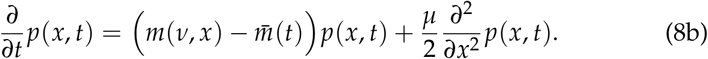

Equation 8b result closely resembles Kimura’s continuum-of-alleles model (Kimura, 1965). The primary difference being that our model utilizes diffusion instead of convolution with an arbitrary mutation kernel. However, our model of mutation can be derived as an approximation to Kimura’s model, which has been referred to as the Gaussian allelic approximation in reference to the distribution of mutational effects on trait values at each locus in a genome (Lande, 1975, Bürger, 1986, Bürger, 2000, Johnson and Barton, 2005), the infinitesimal genetics approximation in reference to modelling continuous traits as being encoded by an infinite number of loci each having infinitesimal effect (Fisher, 1919, Barton et al., 2017) and the Gaussian descendants approximation in reference to offspring trait values being normally distributed around their parental values (Bulmer, 1971, Turelli, 2017).

To distinguish this model from previous models of phenotypic evolution we refer to PDE (4) from which (8b) was derived as the Deterministic Asexual Gaussian allelic model with Abundance dynamics (abbreviated DAGA). Later, we will extend this model to include the effects of demographic stochasticity, which we refer to as the Stochastic Asexual Gaussian allelic model with Abundance dynamics (abbreviated SAGA).

#### Evolutionary Dynamics

We now apply DAGA to derive the dynamics of mean trait 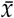 and phenotypic variance *σ*^2^. Both of these dynamics are expressible in terms of covariances with fitness. For an abundance distribution *v*(*x*) and associated phenotypic distribution *p*(*x*), the covariance of fitness and phenotype across the population is defined as

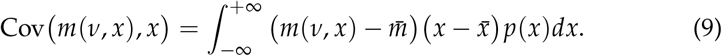

Following this, we again apply integration-by-parts and the chain rule from calculus to find the dynamics of the mean trait 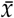 as

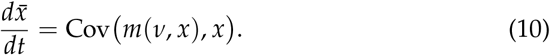

Equation (10) is a continuous time analog of the well known Robertson-Price equation without transmission bias (Frank, 2012, Lion, 2018, Price, 1970, Queller, 2017, Robertson, 1966). Whether or not the covariance of fitness and phenotype creates change in 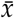 to maximize mean fitness 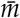 depends on the degree to which selection is frequency dependent (Lande, 1976). Since this change is driven by a covariance with respect to phenotypic diversity, the response in mean trait to selection is mediated by the phenotypic variance. In particular, when *σ*^2^ = 0, 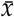 will not respond to selection.

Following the approach taken to calculate the evolution of 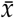, we find the response of phenotypic variation to this model of mutation and selection is

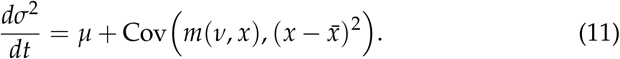

In the absence of mutation equation (11) mirrors the result derived by Lion (2018) for discrete phenotypes. From a statistical perspective, if we think of 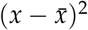 as a square error, then in analogy to the dynamics of the mean trait, we see that the response in *σ*^2^ to selection can be expressed as a covariance of fitness and square error, which is defined in analogy to Cov(*m*(*v, x*),*x*). Just as for the evolution of 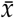, this covariance also creates change in *σ*^2^ that can either increase or decrease mean fitness 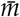, depending on whether or not selection is frequency dependent. The effect of selection on phenotypic variance can be positive or negative depending on whether selection is stabilizing or disruptive.

#### 2.2. Extending DAGA to Demographic Stochasticity

In SM §4, we extend these results to include the effects of demographic stochasticity. The idea is to add an appropriate noise term to DAGA. Hence, we wish to study <u>stochastic</u> partial differential equations (SPDE) that provide natural generalizations of DAGA. Fortunately, rigorous first principle derivations of such SPDE have been provided by Li (1998) and Champagnat et al. (2006). The noise terms driving these SPDE are space-time white noise processes, denoted 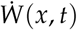, which are random processes uncorrelated in both space and time. In SM §2.1, we provide a set of heuristics for performing calculations with respect to space-time white noise including methods to derive SDE from SPDE in analogy to our derivations of ordinary differential equations (ODE) from PDE above. Since our aim is to present this material to a wide audience of mathematical evolutionary ecologists, our treatment of space-time white noise and stochastic integration deviates from standard definitions to remove the need for a detailed technical treatment. However, in SM §2.2, we show our heuristics are consistent with the rigorous infinite-dimensional stochastic calculus presented in Da Prato and Zabczyk (2014). Using our simplified approach, the reader will only need some elementary probability and an intuitive understanding of SDE, including Brownian motion, in addition to the notions of Riemann integration and partial differentiation already employed.

To understand how SPDE can be derived from biological first principles, we provide in SM §3.1 an informal discussion of measure-valued branching processes (MVBP) (which provide individual-based models) and their diffusion-limits (which provide population-level models). Diffusion-limits of MVBP return so-called superprocesses which track the evolution of abundance and phenotypic distribution (Etheridge, 2000). For univariate traits and under biologically natural conditions, these superprocesses admit abundance densities satisfying SPDE. Under the simplifying assumptions inherited from our treatment of deterministic dynamics and the additional assumption that the variance of individual reproductive output, denoted by *V* ≥ 0, is independent of trait values, we obtain as a special case the relatively simple expression for an SPDE that generalizes DAGA

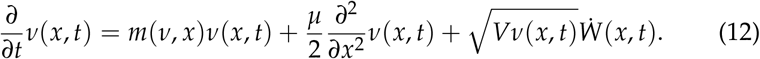

We refer to this special case as the Stochastic Asexual Gaussian allelic model with Abundance dynamics (SAGA). The simplicity of SAGA allows us to use properties of space-time white noise processes to derive a set of SDE that generalize equations (8a), (10) and (11) to include the effects of demographic stochasticity (see SM §3.2 and SM §4). In particular, we find

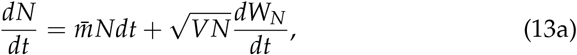

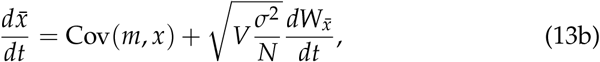

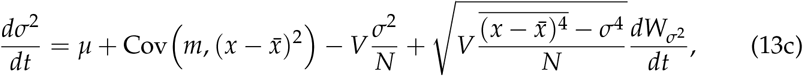

where *W_N_*, 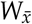 and *W*_*σ*^2^_ are standard Brownian motions and barred expressions such as 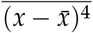 are averaged quantities with respect to the phenotypic distribution *p*(*x, t*). Intuitively, one can interpret equations (13) as if they are ordinary differential equations, but this is not technically rigorous since Brownian motion is nowhere differentiable with respect to time. In SM §4 we show that in general *W_N_* is independent of both 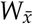 and *W*_*σ*^2^_, but 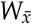 and *W*_*σ*^2^_ may covary depending on the shape of *p*(*x, t*).

Many known results follow directly from expressions (13). Firstly, assuming no variance in reproductive output so that *V* = 0 recovers the deterministic dynamics derived in §2.1. Alternatively, one can take *N* → ∞ to recover the deterministic dynamics for 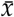 and *σ*^2^. Characteristically, we note the effect of demographic stochasticity on abundance grows with 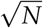. Hence, dividing by *N*, we find the effects of demographic stochasticity on the per-capita growth rate diminish with increased abundance. Relating the response to demographic stochasticity derived here to the effect of random genetic drift derived in classic quantitative genetic theory, if we set *σ*^2^ and *N* constant with respect to time, then integrating the stochastic term in equation (13b) over a single unit of time returns a normally distributed random variable with mean zero and variance equal to *V*_*σ*^2^_/ *N*. In particular, assuming perfect inheritance, when reproductive variance is unity (*V* = 1) this random variable coincides with the effect of random genetic drift on the change in mean trait over a single generation derived using sampling arguments (Lande, 1976). There is also an interesting connection with classical population genetics. A fundamental result from early population genetic theory is the expected reduction in diversity due to the chance loss of alleles in finite populations (Fisher, 1923, Wright, 1931). This expected reduction in diversity due to random genetic drift is captured by the third term in the deterministic component of expression (13c), particularly −*Vσ*^2^/*N*. The component of SDE (13c) describing random fluctuations in *σ*^2^ is more complicated and is proportional to the root of the difference between the centralized fourth moment of the phenotypic distribution and square of the phenotypic variance *σ*^4^.

These expressions can be used to investigate the dynamics of the mean and variance for a very general set of phenotypic distributions. However, in the next subsection we simplify these expressions by assuming normally distributed trait values, known as the Gaussian population assumption (Turelli 2017). In SM §4 we show that under the Gaussian case *W_N_*, 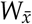 and *W*_*σ*^2^_ are independent. Hence, although the Gaussian population assumption is very restrictive as a model of phenotypic diversity and, except for very special cases of growth rates, is not formally justified, its exceedingly convenient properties make it an important initial approximation.

### 2.3. Particular Results Assuming a Gaussian Phenotypic Distribution

By assuming normally distributed trait values, expressions (13) transform into efficient tools for deriving the dynamics of populations given a fitness function *m*(*v, x*). Gaussian phenotypic distributions can be formally obtained through Gaussian, exponential or weak selection approximations together with a simplified model of mutation, genotype-phenotype mapping and asexual reproduction or random mating (Bürger, 2000, Lande, 1980, Turelli, 2017, 1986, 1984). Hence, given appropriate assumptions on selection, mutation and reproduction, the abundance density *v*(*x, t*) can be approximated as a Gaussian curve in *x* when the ratio *V/N* is small (i.e., when the variance in reproductive output is much smaller than the population size). As with any diffusion approximation, this requires a sufficiently large abundance to accurately reflect the dynamics of populations. Therefore, models developed in this framework are not suitable for studies involving very small population sizes. Allowing for these restrictions, we assume

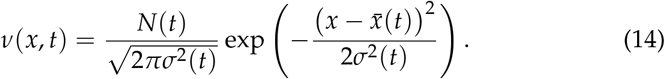

Under this assumption, covariances with fitness can be written in terms of fitness gradients. In particular, we find

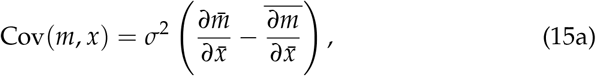

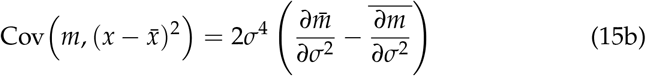

and 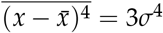. These results imply trait dynamics can be rewritten as

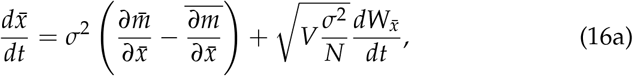

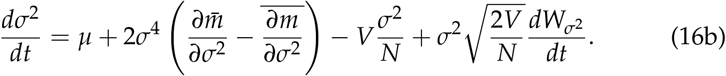

These equations allow us to derive the response in trait mean and variance by taking derivatives of fitness, a much more straightforward operation than calculating a covariance for general phenotypic distributions. Note that in the above expressions, the partial derivatives of 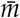 represent frequency independent selection and the averaged partial derivatives of *m* represent frequency dependent selection. This relationship has already been pointed out by Lande (1976) for the evolution of the mean trait in discrete time, but here we see an analogous relationship holds in continuous time and also for the evolution of trait variance.

In SM §5 we generalize this result to the case when traits are imperfectly inherited. In this case, the phenotypic variance *σ*^2^ is replaced by a genetic variance *G*. This genetic variance represents the component of *σ*^2^ explained by additive effects among genetic loci encoding for the focal phenotype (Bulmer, 1971, Roughgarden, 1979, Walsh and Lynch, 2018). It is therefore fitting that *G* is referred to as the additive genetic variance. Following classical quantitative genetic assumptions we find

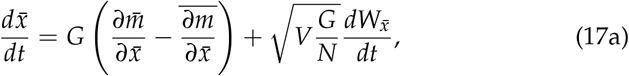

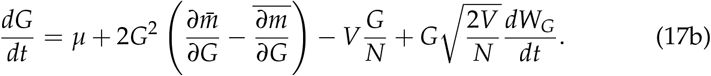

From expressions (17) we see that, under our simple treatment of inheritance, focusing on additive genetic variance *G* instead of the variance in expressed traits *σ*^2^ makes no structural changes to the basic equations describing the dynamics of populations. Instead we see the role played by the variance of expressed traits is now being played by the additive genetic variance. In the next section, we make use of these expressions to develop a model of diffuse coevolution in a guild of *S* species competing along a resource continuum.

## 3. A Model of Diffuse Coevolution

In this section we demonstrate the use of our framework by developing a model of diffuse coevolution across a guild of *S* species whose interactions are mediated by resource competition along a single niche axis. Because our approach treats abundance dynamics and evolutionary dynamics simultaneously, this model allows us to investigate the relationship between selection gradients and competition coefficients, which we carry out in §3.3.

### 3.1. Formulation

The dynamics of phenotypic distributions and abundances have been derived above and so the only task remaining is the formulation of a fitness function. Our approach mirrors closely the theory developed by MacArthur and Levins (1967), Levins (1968) and MacArthur (1972, 1970, 1969). The most significant difference, aside from allowing evolution to occur, is our treatment of resource availability. In particular, we assume resources are replenished rapidly enough to ignore the dynamics of their availability. A derivation from the MVBP framework is provided in SM §6.

#### Abiotic Selection and Competition

For species *i* we inherit the above notation for trait value, distribution, average, variance, abundance, etc., except with an *i* in the subscript. Real world examples of niche axes include the size of seeds consumed by competing finch species and the date of activity in a season for pollinators competing for floral resources. For mathematical convenience, we model the axis of resources by the real line 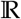. The value of a resource along this axis is denoted by the symbol *ζ*. For an individual in species *i*, we assume resources are sampled from the environment following the utilization curve *u_i_*, which we assume can be written as

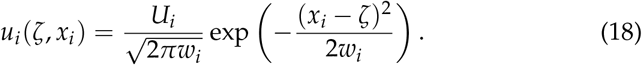

We further assume the niche center *x_i_* is normally distributed among individuals in species *i*, but the niche breadth *w_i_* and total niche utilization *U_i_* are constant across individuals in species *i* and therefore cannot evolve. We assume resources are distributed along the niche gradient and that each species experiences heterogeneous fitness benefits at different niche locations. Taking into account both resource availability and fitness benefits, we suppose individuals of species *i* maximize their benefits by sampling resources at niche location 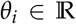. We assume the benefits for individuals of species *i* derived from resources with value 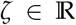 decreases as (*ζ* – *θ_i_*)^2^ increases at a rate *A_i_* ≥ 0. In the absence of competition, we further suppose individuals leave on average *Q_i_* offspring when their utilization curve is concentrated at *θ_i_* (that is, when *x_i_* = *θ_i_* and *w_i_* = 0). Combining these assumptions, we denote by *e_i_*(*ζ*) the fitness benefits for individuals sampling at niche location *ζ* so that

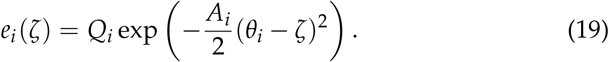

The effect of abiotic stabilizing selection on the fitness for an individual of species *i* with niche location *x_i_* is then given by

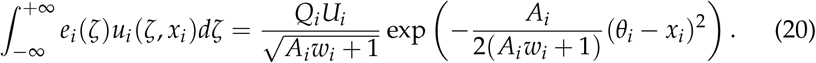

To determine the potential for competition between individuals with niche locations *x_i_* and *x_j_*, belonging to species *i* and *j* respectively, we compute the niche overlap

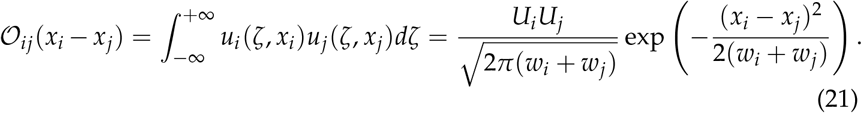

To map the degree of niche overlap to fitness, we assume competition between individuals with niche locations *x_i_* and *x_j_* decreases the expected reproductive output for the individual in species *i* at the rate 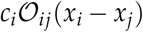 for some *c_i_* > 0. We refer to *c_i_* as the strength of competition for species *i*.

#### The Fitness Function

Assuming the effects due to competitive interactions and abiotic stabilizing selection on the expected reproductive output of individuals accumulates multiplicatively, we derive in SM §6 an expression for the expected reproductive output of individuals in each. Applying a series of diffusion-limits, we then find the following expressions for the growth rate associated with trait value *x* for species *i* along with the population growth rate of species *i*:

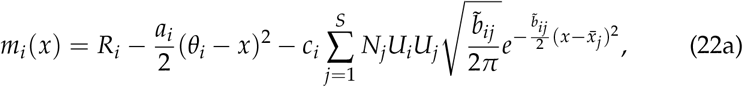

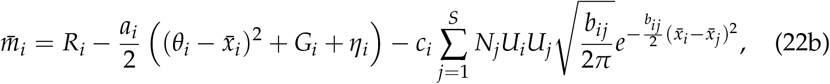

where *a_i_* is the strength of abiotic stabilizing selection on species *i*. The variables 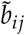, *b_ij_* determine the sensitivity of competitive effects on species *i* to differences in niche locations between species *i* and *j*. We refer to *R_i_* as the innate growth rate of species *i* to distinguish it from the intrinsic growth rate commonly referred to in the field of population ecology. These are composite parameters given by the following expressions:

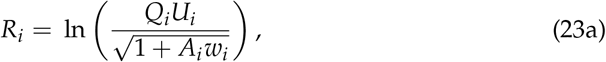

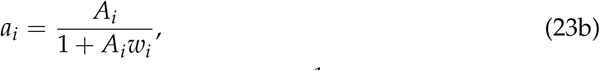

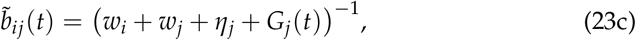

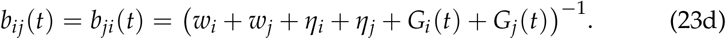

### 3.2. The Model

In SM §6 we combine equations (13a), (17) and (22) to find

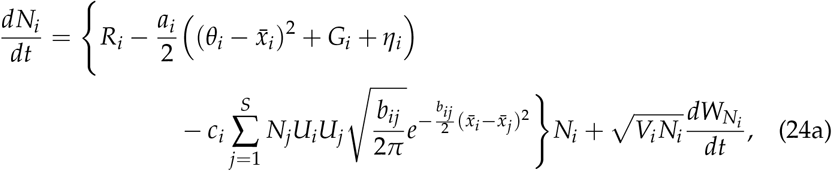

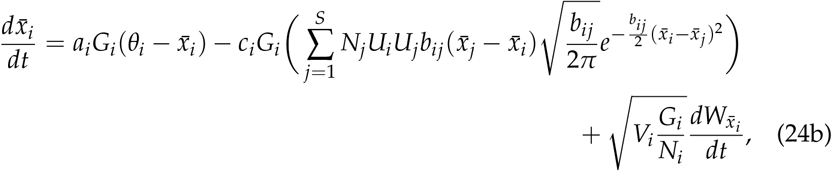

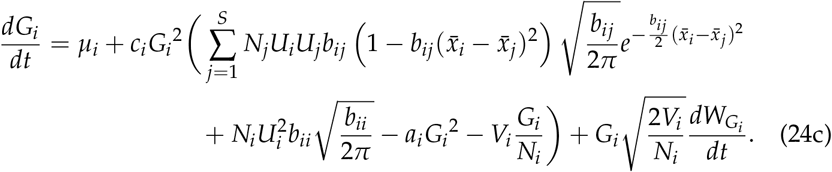

Together, equations (24) provide a synthetic model capturing the dynamics of abundance and evolution from common biological mechanisms.

#### Model Behavior

Despite the convoluted appearance of system (24), there are some nice features that reflect biological reasoning. For example, the dynamics of abundance generalize Lotka-Volterra dynamics. In particular, the effect of competition with species *j* on the fitness of species *i* grows linearly with *N_j_*. However, as biotic selection pushes 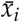 away from 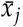, the effect of competition with species *j* on the fitness of species *i* rapidly diminishes due to the Gaussian weights capturing a reduction in niche overlap. These Gaussian weights have been usefully employed to capture interaction preference in recent investigations of coevolution in mutualistic networks (de Andreazzi et al., 2019, Medeiros et al., 2018, Guimarães et al., 2017). The divergence of 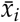 and 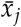 due to competition is referred to in the community ecology literature as character displacement (Brown and Wilson, 1956). We also see that the fitness of species *i* drops quadratically with the difference between 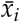 and the abiotic optimum *θ_i_*. Hence, abiotic selection acts to pull 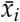 towards *θ_i_*.

The response in mean trait 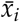 to natural selection is proportional to the amount of heritable variation in the population, represented by the additive genetic variance *G_i_*. However, we have that *G_i_* is itself a dynamic quantity. Under our model, abiotic stabilizing selection erodes away heritable variation at a rate that is independent of both *N_i_* and 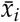. The effect of competition on *G_i_* is a bit more complicated. When 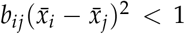, competition with species *j* acts as diversifying selection which tends to increase the amount of heritable variation. However, when 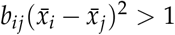, competition with species *j* acts as directional selection and reduces *G_i_*. In the following subsections we demonstrate the behavior of system (24) by plotting numerical solutions and investigate implications for the relationship between the strength of ecological interactions and selection.

#### Community Dynamics

For the sake of illustration we numerically integrated system (24) for a richness of *S* = 100 species. We assumed homogeneous model parameters across species in the community as summarized by Table 1. We repeated numerical integration under the two scenarios of weak and strong competition. For the first scenario of weak competition we set *c* = 1.0 × 10^−7^ and for the second scenario of strong competition we set *c* = 5.0 × 10^−6^. With these two sets of model parameters, we simulated our model for 1000.0 units of time. For both scenarios, we initialized the trait means to 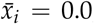, additive genetic variances to *G_i_* = 10.0 and abundances to *N_i_* = 1000.0 for each *i* = 1,…, *S*.

**Table 1:** Values of model parameters used for numerical integration.

| Parameter | Description | Value |
| --- | --- | --- |
| $S$ | species richness | 100 |
| $R$ | innate growth rate, see §3.3 | 1.0 |
| $\theta$ | abiotic optimum | 0.0 |
| $a$ | strength of abiotic selection | 0.01 |
| $c$ | sensitivity to competition | $\{1.0 \times 10^{-7}, 5.0 \times 10^{-6}\}$ |
| $w$ | niche breadth | 0.1 |
| $U$ | total niche use | 1.0 |
| $\eta$ | developmental noise | 1.0 |
| $\mu$ | mutation rate | $1.0 \times 10^{-7}$ |
| $V$ | variance of reproductive output | 5.0 |

Temporal dynamics for each scenario are provided in Figure 1. This figure suggests weaker competition leads to smoother dynamics and a higher degree of organization within the community. Considering expression (24a) we note that, all else equal, relaxed competition allows for larger growth rates which promote greater abundances. From (24a) we also note that the per-capita effects on demographic stochasticity diminish with abundance. To see this, divide both sides by *N_i_*.

**Figure 1:**
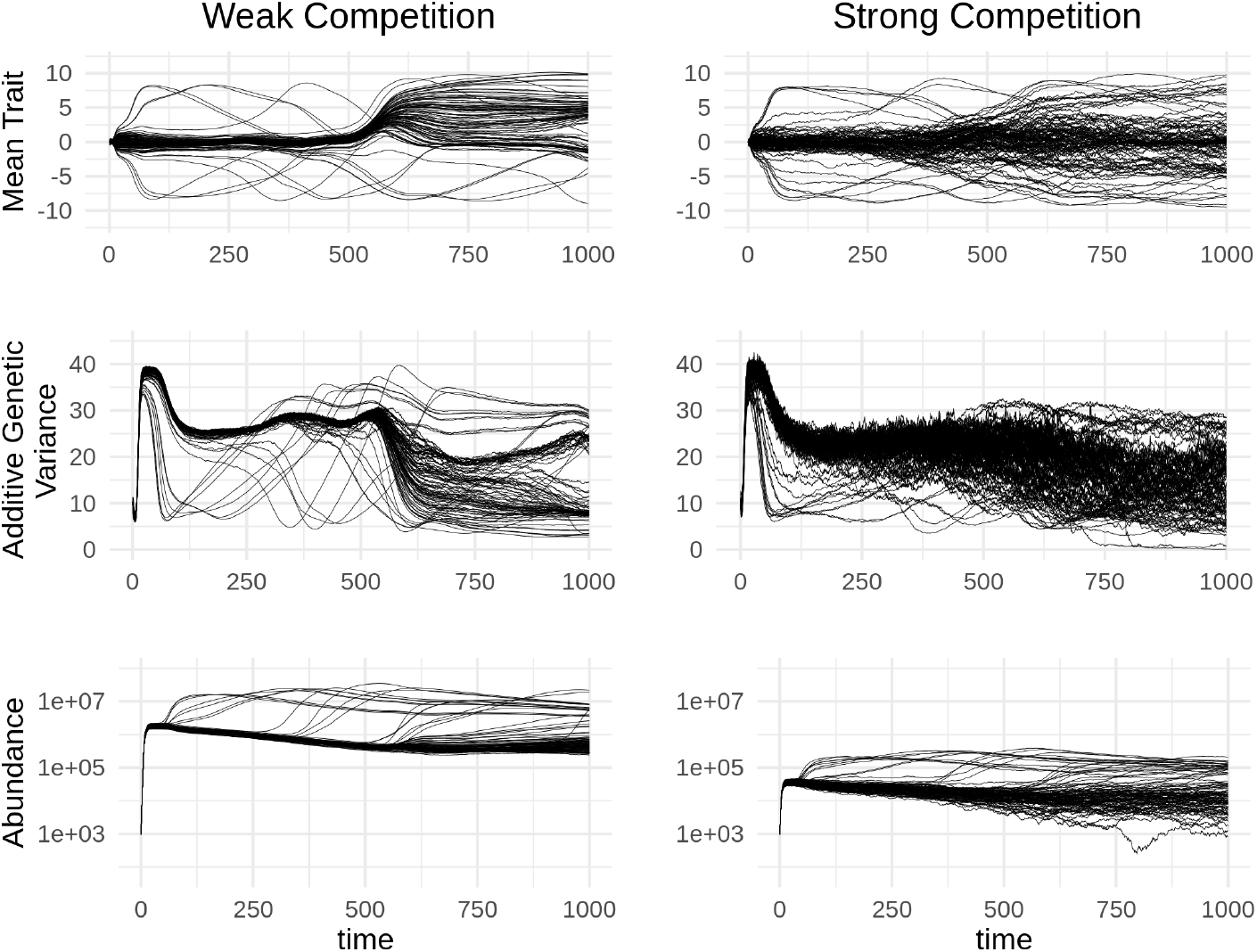
Temporal dynamics of mean trait (top), additive genetic variance (middle) and abundance (bottom) for the scenario of weak competition (left) and strong competition (right).

Inspecting expressions (24b) and (24c), we see that larger abundances also erode the effects of demographic stochasticity on the evolution of mean trait and additive genetic variance. These effects were already noted in §2.2, and thus are not a consequence of our model of coevolution per-se, but we revisit them here since Figure 1 demonstrates the importance of demographic stochasticity in structuring ecological communities even when populations are very large. Hence, contrary to the common assumption that stochastic effects can be ignored for large populations, we find that minute asymmetries generated by demographic stochasticity remain significant drivers of community structure. In particular, although we initialized each species with identical state variables and model parameters, we found an enormous amount of asymmetry in both the evolutionary and abundance dynamics and even some peculiar synchronized shifts. Although future work may show these bizarre features always dissipate after the system has been given sufficient time to evolve, we see demographic stochasticity has pronounced effects on communities experiencing non-equilibrium dynamics.

Although Figure 1 suggests interesting patterns in the dynamics of abundance and trait evolution, a more formal investigation is needed to better understand the relationship between them. In the following subsection we take a step in this direction by approximating correlations between competition coefficients and components of selection gradients induced by interspecific interactions.

### 3.3. The Relation Between the Strength of Ecological Interactions and Selection

Here we investigate the relationship between competition coefficients, which measure the effect of ecological interactions on abundance dynamics, with selection gradients, which measure the magnitude and direction of selection on mean trait and trait variance. We start by considering the expressions of absolute competition coefficients implied by equations (24). However, it turns out absolute competition coefficients display some unfortunate behaviour with respect to our model. We therefore introduce a slightly modified form of absolute competition coefficients. We then provide formula for the components of linear and quadratic selection coefficients corresponding to the effects of interspecific interactions. Lastly, we use a high-richness (large *S*) approximation to determine correlations between competition coefficients and selection gradients across the community. Associated calculations are provided in SM §7.3.

#### Competition coefficients

Relating our treatment of resource competition to theoretical community ecology, the absolute competition coefficient 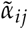, which measures the effect of species *j* on the growth rate of species *i* (sensu Chesson, 2000), becomes a dynamical quantity that can be written as

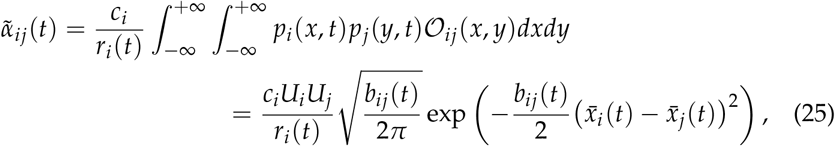

where

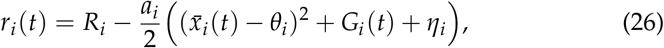

is the intrinsic growth rate of species *i*. Then, *dN_i_*(*t*) can be expressed as

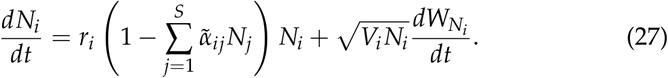

Following our model, the classically defined absolute competition coefficient for species *i* is parameterized with the intrinsic growth rate of species *i* appearing in the denominator. In turn, these intrinsic growth rates depend on the balance between the innate growth rate *R_i_* and the effect of abiotic stabilizing selection. However, this balance further depends on mean trait and additive genetic variance, which evolve freely. This leads to the potential for the signage of *r_i_* to switch between positive and negative which implies the potential for infinite absolute competition coefficients. Furthermore, we see these competition coefficients are influenced by abiotic stabilizing selection instead of solely capturing the effects of inter/intraspecific interactions. Hence, we find it necessary to introduce a modification of the absolute competition coefficient 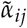 that avoids these caveats. In particular, we define

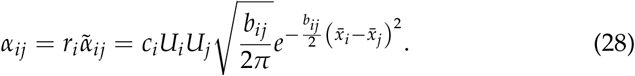

We call *α_ij_* the <u>specific</u> competition coefficient mediating the effects of species *j* on the growth rate of species *i*. Under this parameterization, the abundance dynamics of species *i* is now expressed as

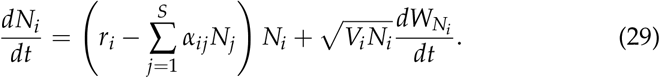

#### Selection Gradients

Linear and quadratic selection gradients have been defined by Lande and Arnold (1983). While the linear selection gradient *β* measures the effect of selection on mean trait evolution, the stabilizing selection gradient *γ* measures the effect of selection on additive genetic or phenotypic variance. Since these quantities are classically defined with respect to discrete-time models of trait evolution, we provide the analogous definitions for continuous-time models in SM §7.1. Following our model of diffuse coevolution, we then show these selection gradients can be additively partitioned into components due to interactions with each species and abiotic stabilizing selection. In particular, we find the components of linear and quadratic selection gradients for species *i* induced by species *j* are given respectively by

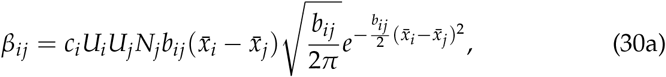

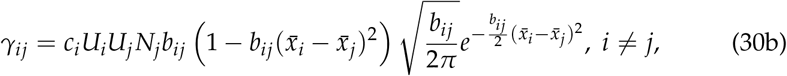

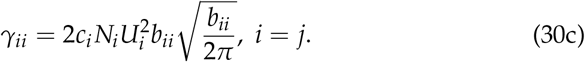

With these expressions, the dynamics of mean trait and additive genetic variance simplify to

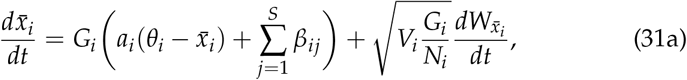

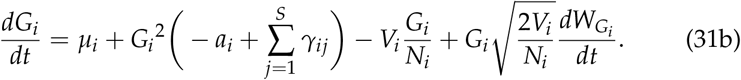

#### High-Richness Approximation

We now make use of the expressions derived for competition coefficients and selection gradients to investigate their relationship. As a first pass, we assume the niche-breadths *w_i_* and intraspecific variances 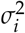 are equivalent across species so that the sensitivity parameters 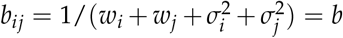 are constant across interacting pairs of species. We also assume abundances *N_i_*, niche-use parameters *U_i_*, strengths of competition *c_i_* and mean traits 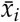 are distributed independently of each other with respective means and variances denoted by 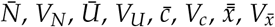. We further assume that richness *S* is large and the distribution of mean trait values is approximately normal.

Under these assumptions we obtained analytical approximations for the correlations between specific competition coefficients *α_ij_* and selection gradients *β_ij_, γ_ij_*. These calculations are provided in SM §7.3. In particular, we found linear selection gradients are not associated with competition coefficients (Corr(*α, β*) ≈ 0). However, we did find a non-trivial relationship between the magnitudes of linear selection gradients and competition coefficients (Corr(*α*, |*β*|) ≠ 0) and also between quadratic selection gradients and competition coefficients (Corr(*α, γ*) ≠ 0). Their expressions can be found in SM §7.3.

To understand if associations between competition coefficients and selection gradients tend to be positive or negative, we visualized these relationships in Figure 2. We fixed 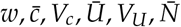 and *V_N_* and allowed the amounts of intraspecific trait variance *σ*^2^ and interspecific trait variance 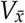 to vary. We found, for biologically realistic areas of parameter space, absolute values of linear selection gradients and quadratic selection gradients tend to be positively associated with competition coefficients. Hence, if we know of competing species that strongly effect each others abundances then we can guess they also impose directional and diversifying selection on one another. However, based on this information alone, we cannot guess at the direction of selection.

**Figure 2:**
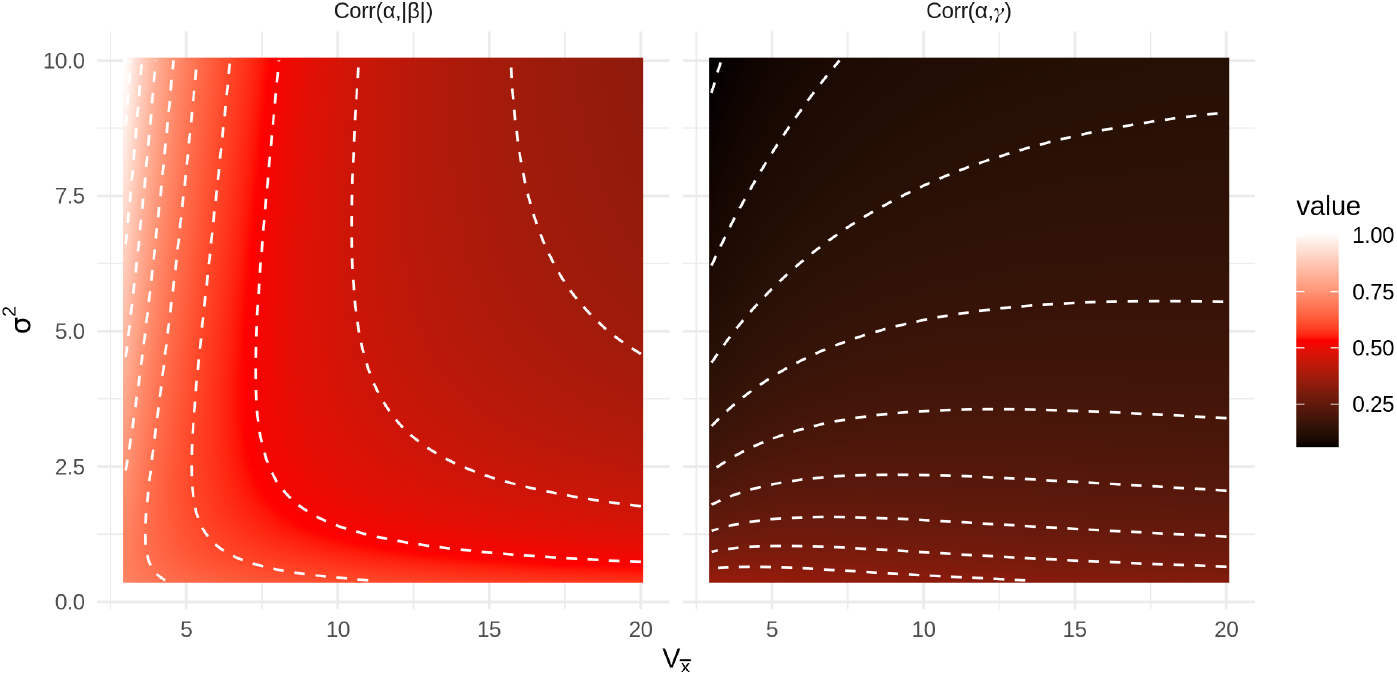
Heatmaps of the correlation between the magnitude of linear selection gradients and competition coefficients (left) and between stabilizing selection gradients and competition coefficients (right) as functions of community-wide variance of mean trait values 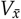 and intraspecific trait variances *σ*^2^. In both plots we set *w* = 1.0, 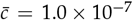, *V_c_* = 0.0, 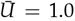, *V_U_* = 0.0, 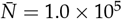, and *V_N_* = 100.0.

## 4. Conclusion

We have introduced a novel approach to derive eco-evolutionary models using the calculus of white noise and diffusion-limits of measure-valued branching processes (MVBP) and coined SAGA, a SPDE model of phenotypic evolution that accounts for demographic stochasticity. From SAGA we derived SDE that track the dynamics of abundance, mean trait and additive genetic variance. Observing the expressions of these SDE, we find the effects of demographic stochasticity on the evolution of mean trait and additive genetic variance characterize the effects of random genetic drift. Although Lande (1976) has previously characterized the effects of random genetic drift on mean trait evolution in quantitative genetic models, the approach taken assumed constant effective population size and discrete non-overlapping generations. In contrast, our approach shows random genetic drift is a result of demographic stochasticity for continuously reproducing populations with fluctuating abundances.

To illustrate the relevance of our approach to studies of evolutionary ecology, we combined our SDE with classical competition theory to derive a model of diffuse coevolution. We then used this model to investigate the relationship between standardized selection gradients and competition coefficients. We found absolute values of linear selection gradients and raw values of quadratic selection gradients are positively related with competition coefficients. In the process, we derived expressions for competition coefficients and components of selection gradients due to pairwise interactions as functions of niche-use parameters (niche breadth, total use and mean and variance of niche location), strength of competitive interactions and abundance.

Although the framework outlined here holds great potential for developing a synthetic theory of coevolving ecological communities, there are two technical gaps in the mathematical foundations of our approach. Firstly, we were unable to derive formal conditions under which trait means and variances remain finite for finite time. However, a result due to Evans and Perkins (1994) shows that the diffusion-limit for a pair of interacting MVBP following our simple niche-based treatment of competition exist when growth rates, as functions of trait values and abundances, are bounded above. This result can be easily extended to finite sets of competing species and therefore formally establishes the existence of abundances as diffusion processes. Further work is needed to determine the conditions under which trait means and variances exist as diffusion processes. The models studied here provide likely sufficient conditions. In particular, since diffusive mutation does not lead to “heavy-tailed” phenotypic distributions, we expect the mean trait and trait variance to remain finite so long as total abundance is positive, given finite initial values for trait mean and variance. That is, since we have not included any processes that would cause blow-up either in mean trait or trait variance, we expect solutions of the SDE (13) to exist for all finite time *t* such that *N*(*t*) > 0 when 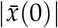, *σ*^2^(0) < +∞. This assumption appears especially well-founded under quadratic stabilizing selection. Since fitness indefinitely decreases as individual trait value becomes indefinitely large (see equation (22)), the diversifying effects of mutation and competition will eventually be overwhelmed by stabilizing selection. Hence quadratic stabilizing selection prevents the abundance densities of populations from venturing indefinitely far from their phenotypic optima.

Secondly, although SDE derived under the assumption of normally distributed phenotypes provide particularly useful formula by replacing covariances between phenotype and fitness with fitness gradients, this assumption is mathematically rigorous only under deterministic dynamics and when the growth rate is a linear or concave-down quadratic function of trait value. However, following our derivation based on classical competition theory, we found the associated growth rate is highly non-linear. While this extreme non-linearity is mathematically inconvenient, it also captures important biological details and thus allows for a more realistic model of community dynamics. In spite of this inconsistency in our model formulation, we found resulting dynamics under the assumption of normally distributed trait values retained well-founded biological intuition. Furthermore, previous work in the field of theoretical quantitative genetics has demonstrated the assumption of normally distributed trait values is robust to fitness functions that select for non-normal trait distributions when inheritance is given a more realistic treatment and when populations reproduce sexually (Turelli and Barton, 1994, Barton et al., 2017). Hence, future work is needed to extend our approach to account for sexual reproduction, more realistic models of inheritance and to investigate the community-level consequences of non-normally distributed trait values.

Overall, this work demonstrates that connecting contemporary theoretical approaches of evolutionary ecology with some fundamental results in the theory of measure-valued branching processes and their diffusion-limits allows for the development of a rigorous, yet flexible approach to synthesizing the dynamics of abundance and distribution of quantitative characters. In particular, equations (13a) and (17) provide a fundamental set of equations for deriving stochastic eco-evolutionary models involving quantitative traits. However, these equations require an expression for growth rates associated with each trait value. Conveniently, equation (SM.34) in SM §3.1 provides a means to derive such growth rates from individual based models. Taken together, these results provide a means to derive analytically tractable dynamics from mechanistic formulations of fitness as a function of phenotype. The derivation of our model of diffuse coevolution, located in SM §6, demonstrates how to derive eco-evolutionary models involving a set of interacting species from biological first principles. Hence, this work provides a novel set of mathematical tools and a tutorial for their use in theoretical studies of evolutionary ecology and therefore paves the way for future work that provides a holistic theoretical treatment of coevolving ecological communities.

