## Supplementary Material for "A White Noise Approach to Evolutionary Ecology"

### 1 Solutions to DAGA

#### 1.1 Sufficient Conditions for Finite Moments Under DAGA

In this section we investigate the conditions under which the trait mean $\bar{x}(t)$, trait variance $\sigma^2(t)$ and abundance $N(t)$ remain finite for finite time $t \geq 0$ when they evolve according to DAGA.

The growth rate expression $m(\nu, x)$ is actually shorthand for the more accurate expression $m\big((K\nu)(x,t), x\big)$ where $K$ is an operator that accounts for nonlocal effects, such as resource competition, on growth rates (Champagnat et al., 2006; Volpert, 2014). In particular, we consider operators of the form $(K\nu)(x,t) = \int_{\mathbb{R}} \kappa(x-y)\nu(y,t)dy$ for some non-negative and bounded function $\kappa$. Hence, $K\nu$ is a non-negative function whenever $\nu$ is a non-negative function. In particular, this implies $m$ is actually a bivariate function of two real numbers $h \geq 0$ and $x \in \mathbb{R}$. To ensure existence and uniqueness of solutions to DAGA, we assume the existence of $R \in \mathbb{R}$ such that $m(h,x) \leq R$ across all $h \geq 0$ and $x \in \mathbb{R}$ along with a twice continuously differentiable and integrable initial condition $u(x)$ that satisfies

$$0 < \int_{\mathbb{R}} (|x| + x^2)u(x)dx < +\infty. \tag{SM.1}$$

In particular, this implies finite initial moments $N(0), |\bar{x}(0)|, \sigma^2(0) < +\infty$ and positive initial abundance and trait variance $0 < N(0), \sigma^2(0)$. Following DAGA, we consider the Cauchy problem

$$\begin{cases} \dot{\nu}(x,t) = m(\nu,x)\nu(x,t) + \frac{\mu}{2}\Delta\nu(x,t) & t > 0 \\ \nu(x,0) = u(x) & t = 0. \end{cases} \tag{SM.2}$$

We assume the operator $F$ defined by $\nu(x,t) \to m(\nu,x)\nu(x,t)$ is locally Lipschitz continuous. This implies that, given two abundance densities $\nu_1(x), \nu_2(x)$ with total abundances $N_1 = \int \nu_1(x)dx$, $N_2 = \int \nu_2(x)dx$ and a positive number $M > 0$, there exists a constant $L_M > 0$ depending on $M$ such that when $N_1, N_2 \leq M$, then

$$\int_{-\infty}^{+\infty} |m(\nu_1,x)\nu_1(x) - m(\nu_2,x)\nu_2(x)|\, dx \leq L_M \int_{-\infty}^{+\infty} |\nu_1(x) - \nu_2(x)| dx. \tag{SM.3}$$

To be specific, we define the domain of the Laplacian as $D(\Delta) = C^2(\mathbb{R}) \cap L^1(\mathbb{R})$ with the norm $\|v\| = \int_{\mathbb{R}} |v(x)|dx$ and define $F$ as an operator on $D(\Delta)$. That is, $F$ maps between functions that are integrable and twice continuously differentiable. Then Theorem 2.5.6 of Zheng (2004) implies for some maximal $T > 0$, the Cauchy problem (SM.2) admits a unique classical solution $\nu(x,t)$ for $t \in [0,T)$. This implies the solution $\nu(x,t)$ is continuously differentiable with respect to $t$ and twice continuously differentiable with respect to $x$ for all $t \in [0,T)$. Furthermore, Theorem 2.5.6 of Zheng (2004) implies either $T = +\infty$ and $N(t) < +\infty$ for all $t > 0$ or $T < +\infty$ and $\lim_{t\uparrow T} N(t) = +\infty$. The latter case corresponds to the notion of "blow-up".

In this section we show that our assumptions on mutation, initial conditions and growth rate, $m(h,x) \leq R$ for all $h \geq 0$ and $x \in \mathbb{R}$ in particular, implies $T = +\infty$ and $N(t) < +\infty$ for all $t > 0$. Replacing $m$ with the upper bound $R \in \mathbb{R}$, PDE (SM.2) reduces to a simple parabolic equation that can be solved using elementary techniques (Farlow, 1993). In particular, when $m(h,x) \equiv R = 0$ we denote the solution to (SM.2) by $\nu_0(x,t)$. Then, denoting

$$\Phi(x,t) = \frac{\exp\left(-x^2/2\mu t\right)}{\sqrt{2\pi\mu t}}, \tag{SM.4}$$

we have

$$\nu_0(x,t) = \int_{\mathbb{R}} \Phi(x-y,t)u(y)dy. \tag{SM.5}$$

In the more general case, when $m(\nu, x) \equiv R \in \mathbb{R}$, equation (SM.2) has the solution $\nu_R(x,t) = e^{Rt}\nu_0(x,t)$. Hence, $\nu_R(x,t) \geq 0$ for all $x \in \mathbb{R}$ and $\int_{\mathbb{R}} \nu_R(x,t)dx = e^{Rt}N(0) < +\infty$ for all $t \geq 0$. Furthermore, denoting

$$N_R(t) = \int_{\mathbb{R}} \nu_R(x,t)dx, \tag{SM.6a}$$

$$p_R(x,t) = \nu_R(x,t)/N_R(t), \tag{SM.6b}$$

$$\bar{x}_R(t) = \int_{\mathbb{R}} x p_R(x,t)dx, \tag{SM.6c}$$

$$\sigma_R^2(t) = \int_{\mathbb{R}} (x - \bar{x}_R(t))^2 p_R(x,t)dx, \tag{SM.6d}$$

we have

$$\bar{x}_R(t) = \int_{\mathbb{R}} x \int_{\mathbb{R}} \Phi(x-y,t)p_R(y,0)dydx = \int_{\mathbb{R}} y p_R(y,0)dy = \bar{x}(0), \tag{SM.7}$$

$$\sigma_R^2(t) = \int_{\mathbb{R}} (x - \bar{x}_R(t))^2 \int_{\mathbb{R}} \Phi(x-y,t)p_R(y,0)dydx = \int_{\mathbb{R}} \Big((y - \bar{x}(0))^2 + \mu t\Big) p_R(y,0)dy = \sigma^2(0) + \mu t. \tag{SM.8}$$

Hence, $|\bar{x}_R(t)|, \sigma_R^2(t) < +\infty$ for all $t \geq 0$. For the sake of contradiction, suppose there exists $x \in \mathbb{R}$ and $t > 0$ such that $\nu(x,t) > \nu_R(x,t)$. Then

$$\nu(x,t) - u(x) = \int_0^t m(\nu,x)\nu(x,s) + \frac{\mu}{2}\Delta\nu(x,s)ds > \int_0^t R\nu_R(x,s) + \frac{\mu}{2}\Delta\nu_R(x,s)ds = \nu_R(x,t) - u(x) \tag{SM.9}$$

which implies there exists $h \geq 0$ and $x \in \mathbb{R}$ such that $m(h,x) > R$. But this contradicts our assumption $m(h,x) \leq R$ for all $h \geq 0$ and $x \in \mathbb{R}$. So we have $\nu(x,t) \leq \nu_R(x,t)$ for each $x \in \mathbb{R}$ and $t \geq 0$. This implies that, for all $t > 0$, $N(t) = \int_{\mathbb{R}} \nu(x,t)dx < +\infty$ and

$$0 \leq \int_{\mathbb{R}} x^2\nu(x,t)dx \leq \int_{\mathbb{R}} x^2\nu_R(x,t)dx < +\infty. \tag{SM.10}$$

Furthermore, since $\nu(x,t)$ is a classical solution of Cauchy problem (SM.2) and since we assumed $N(0) > 0$, we conclude $N(t) > 0$ for all finite $t > 0$. Hence, for each $t > 0$,

$$0 \leq \sigma^2(t) + \bar{x}^2(t) = \frac{1}{N(t)} \int_{\mathbb{R}} x^2\nu(x,t)dx < +\infty. \tag{SM.11}$$

#### 1.2 Equilibrium moments for a population experiencing logistic growth and stabilizing selection under DAGA

Here we show, under DAGA, the population moments $N, \bar{x}$ and $\sigma^2$ evolve to the following stable equilibrium

$$\hat{N} = \tfrac{1}{c}(R - \tfrac{1}{2}\sqrt{a\mu}), \tag{SM.12a}$$

$$\hat{\bar{x}} = \theta, \tag{SM.12b}$$

$$\hat{\sigma}^2 = \sqrt{\tfrac{\mu}{a}}, \tag{SM.12c}$$

given the initial condition $N(0) > 0$ and growth rate

$$m(\nu,x) = R - \frac{a}{2}(\theta - x)^2 - c\int_{\mathbb{R}} \nu(y,t)dy = R - \frac{a}{2}(\theta - x)^2 - cN(t) \tag{SM.13}$$

that satisfies $\theta \in \mathbb{R}$, $a, c, \mu > 0$ and $R > \frac{1}{2}\sqrt{\mu a}$. Following equation (SM.13), mean fitness becomes

$$\bar{m}(t) = R - \frac{a}{2}\left[\left(\theta - \bar{x}(t)\right)^2 + \sigma^2(t)\right] - cN(t), \tag{SM.14}$$

and the ODE for $N(t)$ becomes

$$\frac{d}{dt}N(t) = \left\{R - \frac{a}{2}\left[\left(\theta - \bar{x}(t)\right)^2 + \sigma^2(t)\right] - cN(t)\right\}N(t). \tag{SM.15}$$

Solving for equilibrium total abundance $\hat{N}$ amounts to setting $\frac{d}{dt}N(t) = 0$ and solving for $N(t)$. Ignoring the equilibrium $N(t) = 0$, this reduces to solving $\bar{m}(t) = 0$ for $N(t)$, which, assuming finite equilibrial $\hat{\bar{x}}$ and $\hat{\sigma}^2$, returns

$$\hat{N} = \frac{1}{c}\left\{R - \frac{a}{2}\left[(\theta - \hat{\bar{x}})^2 - \hat{\sigma}^2\right]\right\}. \tag{SM.16}$$

Unfortunately, deriving ODE for $\bar{x}(t)$ and $\sigma^2(t)$ leads to expressions involving higher moments and finding ODE for these higher moments will lead to expressions involving yet even higher moments. To avoid this infinite regression, we find the equilibrium abundance density $\hat{\nu}(x)$ by solving $\frac{\partial}{\partial t}\nu(x,t) = 0$ for $\nu(x,t)$. This implies the following ordinary differential equation

$$\frac{d^2}{dx^2}\hat{\nu}(x) = \left(\frac{2c}{\mu}\hat{N} + \frac{a}{\mu}(\theta - x)^2 - \frac{2R}{\mu}\right)\hat{\nu}(x) \tag{SM.17}$$

which has the solution

$$\hat{\nu}(x) = \frac{\hat{N}}{\sqrt{2\pi}}\left(\frac{a}{\mu}\right)^{\frac{1}{4}}\exp\left(-\sqrt{\frac{a}{\mu}}\frac{(\theta - x)^2}{2}\right). \tag{SM.18}$$

From this expression we infer $\hat{\bar{x}} = \theta$ and $\hat{\sigma}^2 = \sqrt{\frac{\mu}{a}}$. Hence $\hat{N} = \frac{1}{c}\left(R - \frac{1}{2}\sqrt{a\mu}\right)$. To show this equilibrium is stable, we use linear stability analysis. Since $\hat{\nu}(x)$ is Gaussian, we do not run into the same issue with higher moments as above. Furthermore, following equations (16) of the main text, ODE for $\bar{x}(t)$ and $\sigma^2(t)$ can now be expressed as

$$\frac{d}{dt}\bar{x}(t) = \sigma^2(t)\left(\frac{\partial\bar{m}(t)}{\partial\bar{x}(t)} - \overline{\frac{\partial m(t)}{\partial\bar{x}(t)}}\right) = a\sigma^2(t)(\theta - \bar{x}(t)), \tag{SM.19a}$$

$$\frac{d}{dt}\sigma^2(t) = 2\sigma^4(t)\left(\frac{\partial\bar{m}(t)}{\partial\sigma^2(t)} - \overline{\frac{\partial m(t)}{\partial\sigma^2(t)}}\right) + \mu = \mu - a\sigma^4(t). \tag{SM.19b}$$

These expressions confirm our findings that $\hat{\bar{x}} = \theta$ and $\hat{\sigma}^2 = \sqrt{\frac{\mu}{a}}$. Furthermore, calculating

$$\frac{\partial}{\partial\sigma^2(t)}\frac{d}{dt}\sigma^2(t) = -2a\sigma^2(t) \tag{SM.20}$$

and evaluating at $\sigma^2(t) = \hat{\sigma}^2$ demonstrates the equilibrium phenotypic variance is stable when $a, \mu > 0$. Hence, calculating

$$\frac{\partial}{\partial\bar{x}(t)}\frac{d}{dt}\bar{x}(t) = -a\sigma^2(t) \tag{SM.21}$$

and evaluating at $\sigma^2(t) = \hat{\sigma}^2$ and $\bar{x}(t) = \hat{\bar{x}}$ demonstrates the equilibrium phenotypic mean is stable when $a, \mu > 0$. Finally, calculating

$$\frac{\partial}{\partial N(t)} \frac{d}{dt} N(t) = R - \frac{a}{2}\left[\left(\theta - \bar{x}(t)\right)^2 + \sigma^2(t)\right] - 2cN(t) \tag{SM.22}$$

and evaluating at $\sigma^2(t) = \hat{\sigma}^2$, $\bar{x}(t) = \hat{\bar{x}}$ and $N(t) = \hat{N}$ demonstrates the equilibrium total abundance is stable when $a, c, \mu > 0$, and $R > \frac{1}{2}\sqrt{a\mu}$.

### 2 Space-Time White Noise

Here we introduce some heuristics for performing calculations with respect to space-time white noise processes. We then compare these heuristics to results rigorously derived by Da Prato and Zabczyk (2014).

#### 2.1 Heuristics for the White Noise Calculus

We define $\mathfrak{N}_2$ as the set of stochastic processes $f(x,t)$ that are continuous in $t$ and satisfy $\mathbb{E}\left(\int_0^t \int_{\mathbb{R}} |f(x,s)|^2 dxds\right) < +\infty$ for each $t \geq 0$. The operator $\mathbb{E}$ denotes expectation with respect to the underlying probability space. For each $t \geq 0$ we set

$$\|f\|_t = \sqrt{\mathbb{E}\left(\int_0^t \int_{\mathbb{R}} |f(x,s)|^2 dxds\right)}, \tag{SM.23}$$

and make use of the convention $f = g$ if $\|f - g\|_t = 0$ for all $t \geq 0$. Since the abundance density process $\nu(x,t)$ satisfying SAGA is continuous in $t$ and integrable with respect to $x$ for each $t \geq 0$, it also satisfies $\sqrt{\nu} \in \mathfrak{N}_2$. This enables us to utilize the heuristics developed in this section for the derivation of SDE describing the stochastic dynamics of $N(t)$, $\bar{x}(t)$ and $\sigma^2(t)$. To begin developing these heuristics, we introduce a generalized process that captures the essence of space-time white noise in a mathematically tractable format.

We define a generalized stochastic process $\mathbf{W}$ that maps processes $f \in \mathfrak{N}_2$ to real-valued stochastic processes indexed by time $t \geq 0$, but not by space. To evaluate $\mathbf{W}$ for a process $f \in \mathfrak{N}_2$ and some time $t \geq 0$ we write $\mathbf{W}_t(f)$. Specifically, for any $f, g \in \mathfrak{N}_2$, we define $\mathbf{W}(f)$ and $\mathbf{W}(g)$ to be Gaussian processes satisfying, for any $t, t_1, t_2 \geq 0$,

$$\mathbb{E}\big(\mathbf{W}_t(f)\big) = \mathbb{E}\big(\mathbf{W}_t(g)\big) = 0, \tag{SM.24a}$$

$$\mathbb{C}\big(\mathbf{W}_{t_1}(f), \mathbf{W}_{t_2}(g)\big) = \mathbb{E}\left(\int_0^{t_1 \wedge t_2} \int_{\mathbb{R}} f(x,s)g(x,s)dxds\right), \tag{SM.24b}$$

where $t_1 \wedge t_2 = \min(t_1, t_2)$ and $\mathbb{C}$ denotes covariance with respect to the underlying probability space. In particular, denoting $\mathbb{V}$ the variance operator with respect to the underlying probability space, we have $\mathbb{V}\big(\mathbf{W}_t(f)\big) = \|f\|_t^2$ for all $t \geq 0$ and $f \in \mathfrak{N}_2$.

The operators $\mathbb{E}$ and $\mathbb{C}$ are to be distinguished from expectations and covariances with respect to phenotypic diversity such as $\bar{x}$ and $\mathrm{Cov}(m,x)$. In particular, since we model phenotypic diversity as a random process, the phenotypic moments $\bar{x}$ and $\mathrm{Cov}(m,x)$ are random variables and $\mathbb{E}(\bar{x})$, $\mathbb{E}\big(\mathrm{Cov}(m,x)\big)$ denote the expectations of these random variables with respect to the underlying probability space.

Since Gaussian processes are characterized by their expectations and covariances and since we assume the $\mathfrak{N}_2$ processes are continuous in time, the processes $\mathbf{W}(f)$ and $\mathbf{W}(g)$ are well defined. As an example, if $f \in \mathfrak{N}_2$ is independent of time, then $\mathbf{W}(f)$ is a Brownian motion with variance at time $t \geq 0$ equal to $\|f\|_t^2 = t\,\mathbb{E}(\int_{\mathbb{R}} f^2(x,0)dx)$. With the generalized process $\mathbf{W}$ defined, we define the space-time white noise $\dot{W}(x,t)$ implicitly via the stochastic integral

$$\text{``}\int_0^t \int_{\mathbb{R}} f(x,s)\dot{W}(x,s)dxds\text{''} = \mathbf{W}_t(f),\ \forall\, f \in \mathfrak{N}_2,\ t \geq 0. \tag{SM.25}$$

We place quotations in the above expression to emphasize its informal nature and that it should not be confused with classical Riemann integration. Using this informal notation, equations (SM.24a) and (SM.24b) can be rewritten as

$$\mathbb{E}\left(\int_0^t \int_{\mathbb{R}} f(x,s)\dot{W}(x,s)dxds\right) = 0, \tag{SM.26a}$$

$$\mathbb{C}\left(\int_0^{t_1} \int_{\mathbb{R}} f(x,s)\dot{W}(x,s)dxds, \int_0^{t_2} \int_{\mathbb{R}} g(x,s)\dot{W}(x,s)dxds\right) = \int_0^{t_1 \wedge t_2} \int_{\mathbb{R}} f(x,s)g(x,s)dxds. \tag{SM.26b}$$

To relate these formula to the common notation used for SDE, we write

$$\hat{f}(x,t) = \frac{f(x,t)}{\sqrt{\int_{\mathbb{R}} f^2(y,t)dy}} \quad \text{and} \quad d\hat{\mathbf{W}}_t(f) = \left(\int_{\mathbb{R}} \hat{f}(x,t)\dot{W}(x,t)dx\right) dt \tag{SM.27}$$

so that

$$\int_0^t d\hat{\mathbf{W}}_s(f) = \int_0^t \int_{\mathbb{R}} \frac{f(x,s)}{\sqrt{\int_{\mathbb{R}} f^2(s,y)dy}} \dot{W}(x,s)dxds. \tag{SM.28}$$

This implies

$$\mathbb{E}\left(\int_0^t d\hat{\mathbf{W}}_s(f)\right) = 0,\ \mathbb{C}\left(\int_0^{t_1} d\hat{\mathbf{W}}_s(f), \int_0^{t_2} d\hat{\mathbf{W}}_s(f)\right) = t_1 \wedge t_2 \tag{SM.29}$$

and in particular, as a function of $t$, $\int_0^t d\hat{\mathbf{W}}_s(f)$ is a standard Brownian motion for any $f \in \mathfrak{N}_2$. Hence, $d\hat{\mathbf{W}}_t(f)$ is analogous to the traditional shorthand used to denote stochastic differentials. Thus, equation (SM.26b) effectively extends Itô's multiplication table to Table S1.

Table S1: An extension of Itô's multiplication table.

| $\times$ | $d\hat{\mathbf{W}}_t(f)$ | $d\hat{\mathbf{W}}_t(g)$ | $dt$ |
|---|---|---|---|
| $d\hat{\mathbf{W}}_t(f)$ | $dt$ | $\left(\int_{\mathbb{R}} \hat{f}(x,t)\hat{g}(x,t)dx\right) dt$ | 0 |
| $d\hat{\mathbf{W}}_t(g)$ | $\left(\int_{\mathbb{R}} \hat{f}(x,t)\hat{g}(x,t)dx\right) dt$ | $dt$ | 0 |
| $dt$ | 0 | 0 | 0 |

The extension of Itô's multiplication table and properties of white noise outlined in this subsection provide a useful set of tools for working with SPDE. In SM §4 we employ these tools to derive SDE that track the dynamics of abundance, mean trait and phenotypic variance of a population from a particular SPDE. In the following subsection, we review how this particular SPDE naturally arises as the diffusion limit of a measure-valued branching process (MVBP).

#### 2.2 Comparing the White Noise Heuristics to the Infinite-Dimensional Stochastic Calculus of Da Prato and Zabczyk (2014)

Our above approach is inspired by the treatment provided in §4.2 of Da Prato and Zabczyk (2014). Here the authors develop a stochastic integral of operator-valued processes. In particular, they consider processes indexed by time $t \geq 0$ valued as Hilbert-Schmidt operators $\Phi(t)$ and define the norm

$$\|\Phi\|_t = \sqrt{\mathbb{E}\left(\int_0^t \mathrm{Tr}[\Phi(s)\Phi^*(s)]ds\right)},\ t \geq 0. \tag{SM.30}$$

In our case we only consider the so-called multiplication operators. That is, processes that consist of operators $\Phi(t)$ having the form $\Phi(t)g(x) = \varphi(x,t)g(x)$ such that $\varphi(\cdot,t) \in L^2(\mathbb{R})$ a.s. for each $t \geq 0$. In this case $\Phi(t) = \Phi^*(t)$ and

$$\|\Phi\|_t = \|\varphi\|_t = \sqrt{\mathbb{E}\left(\int_0^t \int_{\mathbb{R}} \varphi^2(x,s)dxds\right)},\ t \geq 0. \tag{SM.31}$$

Da Prato and Zabczyk (2014) form the space $\mathfrak{N}^2_W(0,T)$ of Hilbert-Schmidt operator-valued predictable processes $\Phi(t)$ that satisfy $\|\Phi\|_T < +\infty$ for some $T > 0$. This corresponds to our more specialized space $\mathfrak{N}_2$ that consists of $L^2(\mathbb{R})$-valued processes $\varphi(x,t)$ such that $\|\varphi\|_t < +\infty$ for all $t \geq 0$. In their treatment, $W(t)$ plays a similar role to our generalized process $\mathbf{W}_t$. For $\Phi \in \mathfrak{N}^2_W(0,T)$, they denote the stochastic integral for $t \in [0,T]$ by $\Phi \cdot W(t)$. Hence, for $\Phi(t)g(x) = \varphi(x,t)g(x)$ as above, $\mathbf{W}_t(\varphi) = \Phi \cdot W(t)$. The authors then prove the following:

**Proposition 4.28** *Assume that* $\Phi_1, \Phi_2 \in \mathfrak{N}^2_W(0,T)$. *Then*

$$\mathbb{E}(\Phi_i \cdot W(t)) = 0,\ \ \mathbb{E}(\|\Phi_i \cdot W(t)\|^2) < +\infty,\ \ \forall\ t \in [0,T].$$

**Corollory 4.29** *Under the same assumptions as Proposition 4.28,*

$$\mathbb{C}(\Phi_1 \cdot W(t), \Phi_2 \cdot W(s)) = \mathbb{E}\left(\int_0^{t\wedge s} \mathrm{Tr}[\Phi_2(r)\Phi_1^*(r)]dr\right),\ \ \forall\ t,s \in [0,T].$$

Simplifying these expressions for the multiplication operators described above returns equations (SM.26a) and (SM.26b) above.

### 3 Measure-Valued Branching Processes

#### 3.1 From Branching Processes to SPDE

To represent measure-valued branching processes (MVBP), we denote by $\delta_{x_i(t)}$ the point-mass centered on $x_i(t) \in \mathbb{R}$ representing the $i$-th individual at time $t$. Then the whole population can be represented by

$$X(t) = \sum_{i=1}^{n(t)} \delta_{x_i(t)}. \tag{SM.32}$$

We write $X(D,t)$ to calculate the mass of the population located in $D \subset \mathbb{R}$ at time $t$. Individuals branch between exponentially distributed intervals of time at the rate $\lambda$. For the sake of simplicity we assume the parent dies at a branching event and leaves a random, possibly zero, number of offspring at its final "position". Although it is tradition to denote the lifetime reproductive output (i.e., fitness) of individuals by $W$, this symbol has been taken by Brownian motion. We therefore denote $\mathfrak{W}$ and $V$ respectively the mean and variance of offspring produced at a branching event. When fitness depends on phenotype, our assumptions imply that the parental trait $x_i(t)$ is evaluated at the time $t$ for which parent dies. Mutation is captured by the "spatial movement" of $x_i(t)$, which we assume follows a Brownian motion with diffusion parameter $\sqrt{\mu}$. Hence, offspring trait values will be normally distributed around their parental trait value, which implies the Gaussian descendants approximation coined by Turelli (2017).

To obtain a diffusion-limit of the MVBP $X(t)$, which we refer to as a superprocess (sensu Etheridge, 2000), we consider the initial condition $X(0)$ with initial number of individuals $n(0) = n_0$. We rescale individual mass by $N_0/n_0$, where $N_0$ is any positive number, branching rate by $\lambda \to n_0$, and fitness by $\mathfrak{W} \to \mathfrak{W}^{1/n_0}$ and take the limit $n_0 \to +\infty$. Denoting this rescaling of $X(t)$ by $X^{(n_0)}(t)$, we have

$$X^{(n_0)}(0) = \frac{N_0}{n_0} \sum_{i=1}^{n_0} \delta_{x_i(0)}. \tag{SM.33}$$

From this we see the initial total mass is $N_0$ for each $n_0 = 1, 2, \ldots$. In the diffusion-limit, the particle view of the population is replaced by blob spread across phenotypic space (see Figure S1 for the analogous rescaling of a branching random walk). When $\mathfrak{W} = 1$, this stochastic blob is referred to as a Dawson-Watanabe superprocess.

The existence of diffusion limits for a large class of MVBP allowing for individual interactions has been treated by Méléard and Roelly (1992, 1993). The interactions can manifest as dependencies of the spatial movement or reproductive law of individuals on their position and the state of the whole population. An important result of Méléard and Roelly (1992, 1993) is a formula that provides a method to calculate the continuous-time growth rate for the diffusion-limit $\mathfrak{X}$ from the fitness function $\mathfrak{W}(X, x)$. Following our above rescaling, we have

$$m(\mathfrak{X}, x) = \lim_{n_0 \to \infty} n_0 \left( \mathfrak{W}^{1/n_0}(X^{(n_0)}, x) - 1 \right). \tag{SM.34}$$

Li (1998) built directly off of the construction of Méléard and Roelly (1992, 1993) to study properties of interacting superprocesses and, by assuming individual spatial movement occurs independently of location $x$ and the entire population $X(t)$, showed the evolution of associated density processes can be described by a SPDE. Assuming the interactions manifest only in the fitness function $\mathfrak{W}(X, x)$ and that the growth-rate $m(\mathfrak{X}, x)$ is bounded above and below, Li's (1998) result implies the interacting superprocess on one dimensional trait space has a density $\nu(x, t)$ which is non-negative, integrable, continuous in time and space and satisfies the SPDE

$$\frac{\partial}{\partial t}\nu(x,t) = m(\nu, x)\nu(x,t) + \frac{\mu}{2}\frac{\partial^2}{\partial x^2}\nu(x,t) + \sqrt{V\nu(x,t)}\dot{W}(x,t), \tag{SM.35}$$

where $m(\nu, x) = m(\mathfrak{X}, x)$. We refer to SPDE (SM.35) as the Stochastic Asexual Gaussian allelic model with Abundance dynamics (abbreviated SAGA).

By assuming growth rates are only bounded above, Evans and Perkins (1994) proved a result that demonstrates the existence and uniqueness for multiple interacting superprocesses. The result is known as a bivariate Girsanov transform and formally demonstrates existence for a pair of interacting superprocesses engaged in resource competition following our treatment provided in SM §6. By lumping the pair of interacting superprocesses into a single superprocess such that individuals are now represented by a discrete trait indicating which species they belong to in addition to their trait value, we can consider interactions with yet another superprocess and in this way extend the bivariate Girsanov transform to a multivariate Girsanov transform which then establishes existence of $S$ superprocesses engaged in resource competition. Although we are unaware of conditions under which these superprocesses admit density processes that satisfy SPDE, the derivations of SDE from SPDE using weak solutions given below can be thought of as shorthand for deriving SDE from superprocesses by extending the dual space to include the test functions $f(x) = 1, x, x^2$ when possible. We therefore continue our treatment from the SPDE perspective in what follows.

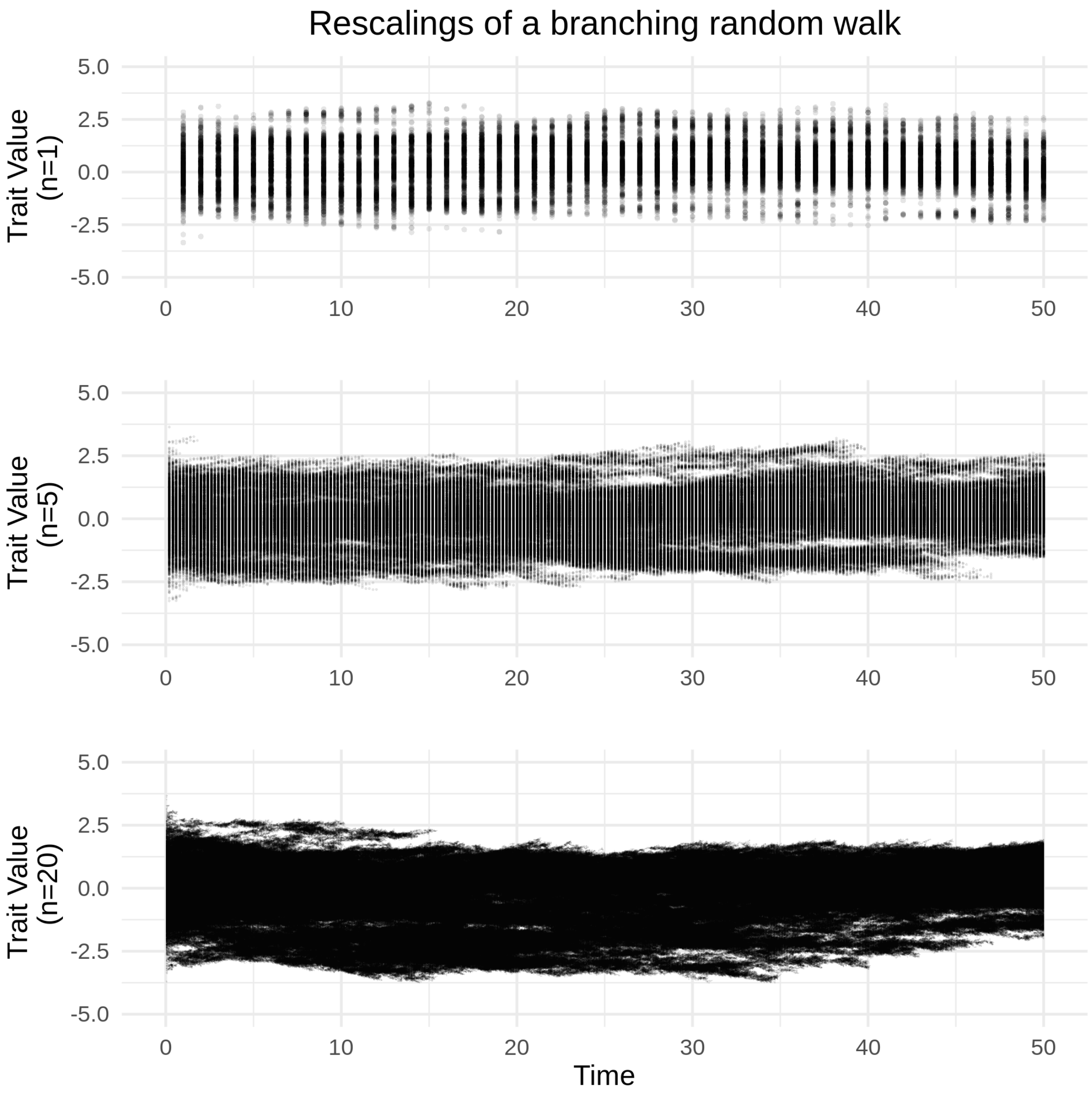

Figure S1: Rescaled sample paths of a branching random walk under stabilizing selection and logistic growth. The top plot displays a sample path without scaling $(n_0 = 1)$, the middle plot shows a sample path rescaled by $n_0 = 5$ and the bottom plot shows a sample path rescaled by $n_0 = 20$. For each of these plots we have set the innate growth rate to $R = 1.0$, phenotypic optimum to $\theta = 0$, strength of abiotic stabilizing selection to $a = 0.01$, sensitivity to competition to $c = 2 \times 10^{-3}$ and mutation rate to $\mu = 1 \times 10^{-3}$.

#### 3.2 Deriving SDE From SPDE

Assuming a growth rate $m(\nu, x)$ such that solutions to SAGA are well defined, we can calculate the total mass process $N(t)$ using the weak solution of SPDE (SM.35) with $f(x) \equiv 1$ (Walsh, 1986; Etheridge, 2000; Evans, 2010). This implies

$$\begin{aligned} N(t) - N(0) &= \int_0^t \int_{\mathbb{R}} \nu(x,s) \left( m(\nu, x) \cdot 1 + \frac{\mu}{2} \frac{\partial^2}{\partial x^2} 1 \right) + 1 \sqrt{V \nu(x,s)} \dot{W}(x,s) ds dx \\ &= \int_0^t \bar{m}(s) N(s) dt + \sqrt{V} \int_0^t \sqrt{N(s)} d\hat{\mathbf{W}}_s \left( \sqrt{\nu(x,s)} \right), \end{aligned} \quad \text{(SM.36)}$$

where the population growth rate is calculated as

$$\bar{m}(t) = \frac{1}{N(t)} \int_{\mathbb{R}} m(\nu, x) \nu(x,t) dx, \quad \text{(SM.37)}$$

and $\hat{\mathbf{W}}_s \left( \sqrt{\nu(x,s)} \right)$ is a standard Brownian motion (see SM §2.1) given by

$$\int_0^t d\hat{\mathbf{W}}_s \left( \sqrt{\nu(x,s)} \right) = \int_0^t \int_{\mathbb{R}} \frac{\sqrt{\nu(x,s)}}{\sqrt{\int_{\mathbb{R}} \nu(x,s) dx}} \dot{W}(x,s) dx ds. \quad \text{(SM.38)}$$

Setting $W_N(t) = \hat{\mathbf{W}}_t(\sqrt{\nu(x,t)})$, we can use traditional stochastic differential notation to write

$$dN = \bar{m} N dt + \sqrt{VN} dW_N. \quad \text{(SM.39)}$$

To find the associated SDE for $\bar{x}(t)$ and $\sigma^2(t)$, we want to repeat the same approach for $f(x) = x, x^2$ and apply Itô's lemma. However, for these cases $f \notin C_b^2(\mathbb{R})$ since $f$ will not be bounded. But, if we can show $\int_{\mathbb{R}} (|x| + x^2 + x^4) \nu(x,t) dx < +\infty$ for all $t > 0$ given this condition is satisfied by $\nu(x, 0)$, then we can apply the weak solution of (SM.35) to derive SDE for $\bar{x}(t)$ and $\sigma^2(t)$. To illustrate, let us suppose this is the case. Setting $\tilde{x}(t) = \int_{\mathbb{R}} x \nu(x,t) dx$, we have

$$\tilde{x}(t) = \tilde{x}(0) + \int_0^t \int_{\mathbb{R}} \nu(x,s) m(\nu, x) x + x \sqrt{V \nu(x,s)} \dot{W}(x,s) dx ds. \quad \text{(SM.40)}$$

Similarly, setting $\tilde{\sigma}^2(t) = \int_{\mathbb{R}} x^2 \nu(x,t) dx$, we have

$$\tilde{\sigma}^2(t) = \tilde{\sigma}^2(0) + \int_0^t \int_{\mathbb{R}} \nu(x,s) \left( m(\nu, x) x^2 + \mu \right) + x^2 \sqrt{V \nu(x,s)} \dot{W}(x,s) dx ds. \quad \text{(SM.41)}$$

Since $\bar{x}(t) = \tilde{x}(t)/N(t)$ and $\sigma^2(t) = \tilde{\sigma}^2(t)/N(t) - \bar{x}^2(t)$, we can use Itô's lemma to derive SDE for $\bar{x}(t)$ and $\sigma^2(t)$, which we perform in SM §4. We make no attempt in finding sufficient conditions to ensure $\int_{\mathbb{R}} (|x| + x^2 + x^4) \nu(x,t) dx < +\infty$ and hence make no general assertions about the existence or uniqueness of $\bar{x}(t)$ or $\sigma^2(t)$. Regardless, we will later assume $\nu(x,t)$ can be approximated by a Gaussian curve in $x$ for all $t \geq 0$. This assumption implies $\int_{\mathbb{R}} |x|^n \nu(x,t) dx < +\infty$ for all $n \in \{1, 2, \dots\}$ and for all $t \geq 0$ and hence guarantees the existence of $\bar{x}(t)$ and $\sigma^2(t)$ for all $t > 0$.

#### 3.3 Simulating the Rescaled Process

Here we provide a detailed description of the branching random walk and how we have chosen to rescale it. To reduce the potential for extinction and to keep the population density concentrated near a particular trait value, we focus on the case of logistic growth and stabilizing selection as described in equation (SM.13). In particular, we focus on a growth rate which, as a function of trait value $x \in \mathbb{R}$ and a measure-valued process $X$, can be written as

$$m(X(t), x) = R - \frac{a}{2}(\theta - x)^2 - cX(\mathbb{R}, t), \tag{SM.42}$$

where $X(\mathbb{R}, t)$ is the total mass of the population at time $t$. We have implemented this simulation in the programming language Julia. A copy can be found at the url:

`https://github.com/bobweek/branching.brownian.motion.and.spde`

##### 3.3.1 Description of Simulation

We begin by describing the branching random walk before introducing our scheme to rescale it. Our branching random walk follows closely the description of measure-valued branching processes in SM §3.1. However, we replace exponentially distributed lifetimes with deterministic unit time steps for easier implementation. Hence, we restrict time to $t = 0, 1, 2, \ldots$, and so on. Furthermore, we allow individual fitness to depend on both trait value and the state of the entire population. At time $t$ we write $\{x_1(t), \ldots, x_{n(t)}(t)\}$ as the set of trait values across all $n(t)$ individuals alive in the population. Since our simulation follows discrete individuals instead of continuous distributions of trait values, we can write

$$X(t) = \sum_{i=1}^{n(t)} \delta_{x_i(t)}, \tag{SM.43}$$

where $\delta_{x_i(t)}$ denotes the point-mass located at $x_i(t)$. For simplicity we assume perfect heritability. At each iteration we draw, for each individual, a random number of offspring from a Negative-Binomial distribution. We use the Negative-Binomial distribution so that we can fix the variance in reproductive output while allowing the mean reproductive output to change. In particular, this coincides with our treatment of diffusion limits of interacting measure-valued processes above.

Recall the Negative-Binomial distribution models the number of failed Bernoulli trials that occur before a given number of successful trials. Denoting $q$ the probability of success for each trial and $s$ the number of successes, the mean and variance are given respectively by

$$\mathfrak{W} = \frac{s(1-q)}{q}, \ V = \frac{s(1-q)}{q^2}. \tag{SM.44}$$

This imposes the restriction $V > \mathfrak{W}$. Requiring the $i$th individual to have mean number offspring $\mathfrak{W}(X, x_i)$ and variance equal to $V$, the parameters of the associated Negative-Binomial distribution become

$$q(X, x_i) = \frac{\mathfrak{W}(X, x_i)}{V}, \ s(X, x_i) = \frac{\mathfrak{W}^2(X, x_i)}{V - \mathfrak{W}(X, x_i)}. \tag{SM.45}$$

For each offspring produced by the individual with trait value $x_i$, we assign independently drawn trait values normally distributed around $x_i$ with variance $\mu$. This summarizes the basic structure of our simulation. To impose selection and density dependent growth rates, we set

$$\mathfrak{W}(X, x_i) = \exp\left(R - \frac{a}{2}(\theta - x_i)^2 - cX(\mathbb{R}, t)\right), \tag{SM.46}$$

where $X(\mathbb{R}, t) = n(t)$.

##### 3.3.2 Rescaling

To rescale the branching random walk by a positive integer $n_0$, we replace individual mass with $\frac{N_0}{n_0}$ for some fixed continuously varying number $N_0 > 0$, write the rescaled population distribution as $X^{(n_0)}$, rescale generation time by $1/n_0$ (which implies mutational variance is rescaled by $1/n_0$) and expected reproductive output by

$$\begin{aligned}\mathfrak{W}(X^{(n_0)}, x_i) \to \left(\mathfrak{W}(X^{(n_0)}, x_i)\right)^{1/n_0} &= \exp\left(\frac{R}{n_0} - \frac{a}{2n_0}(\theta - x_i)^2 - \frac{c}{n_0^2}X^{(n_0)}(\mathbb{R}, \cdot)\right) \\ &= \exp\left(\frac{R}{n_0} - \frac{a}{2n_0}(\theta - x_i)^2 - \frac{cn}{n_0^2}\right). \quad \text{(SM.47)}\end{aligned}$$

When it exists, we denote by $\mathfrak{X} = \lim_{n_0 \to \infty} X^{(n_0)}$, the limiting process of the sequence of rescaled processes $X^{(1)}, X^{(2)}, \ldots$. Then, as desired, we find

$$\lim_{n_0 \to \infty} n_0 \left(\left(\mathfrak{W}(X^{(n_0)}(t), x)\right)^{1/n_0} - 1\right) = R - \frac{a}{2}(\theta - x)^2 - c\mathfrak{X}(\mathbb{R}, t). \quad \text{(SM.48)}$$

Since the limiting growth rate is bounded above by $R$, the bivariate Girsanov transform given by Evans and Perkins (1994) can be used to demonstrate existence and uniqueness of $\mathfrak{X}$ (see also §7.5 of Etheridge, 2000).

### 4 Derivation of SDE for $\bar{x}$ and $\sigma^2$

Here we derive the stochastic dynamics of $\bar{x}$ and $\sigma^2$ under SAGA by combining weak solutions of SPDE, an extension Itô's multiplication table summarized in Table S1 and Itô's quotient rule. This calculation requires the abundance density $\nu(x,t)$ to have finite first, second and fourth phenotypic moments. Hence, we assume

$$\int_{\mathbb{R}} \nu(x,t)(|x| + x^2 + x^4)dx < +\infty. \quad \text{(SM.49)}$$

Following SM §3.1, we set

$$\tilde{x}(t) = \int_{\mathbb{R}} x\nu(x,t)dx, \;\; \tilde{\sigma}^2(t) = \int_{\mathbb{R}} x^2\nu(x,t)dx. \quad \text{(SM.50)}$$

Applying the weak solution of SAGA we obtain diffusion processes defined by

$$N(t) = N(0) + \int_0^t \int_{\mathbb{R}} \nu(x,s)m(\nu,x) + x\sqrt{V\nu(x,s)}\dot{W}(x,s)dxds, \quad \text{(SM.51a)}$$

$$\tilde{x}(t) = \tilde{x}(0) + \int_0^t \int_{\mathbb{R}} \nu(x,s)m(\nu,x)x + x\sqrt{V\nu(x,s)}\dot{W}(x,s)dxds, \quad \text{(SM.51b)}$$

$$\tilde{\sigma}^2(t) = \tilde{\sigma}^2(0) + \int_0^t \int_{\mathbb{R}} \nu(x,s)\left(m(\nu,x)x^2 + \mu\right) + x^2\sqrt{V\nu(x,s)}\dot{W}(x,s)dxds. \quad \text{(SM.51c)}$$

In the following two sections we use Itô's quotient rule to derive expressions for the evolution of $\bar{x} = \tilde{x}/N$ and $\sigma^2 = \tilde{\sigma}^2 - \bar{x}^2$. Following these two sections we investigate stochastic dependencies between the processes $N$, $\bar{x}$ and $\sigma^2$.

#### 4.1 Derivation for Trait Mean

We make use of the notation

$$\begin{cases} \|N\|_2 = \sqrt{V \int_{\mathbb{R}} \nu(x,t)dx} = \sqrt{VN} \\ \|\tilde{x}\|_2 = \sqrt{V \int_{\mathbb{R}} x^2 \nu(x,t)dx} \\ \langle \tilde{x}, N \rangle = V \int_{\mathbb{R}} x\nu(x,t)dx = \bar{x}VN. \end{cases} \tag{SM.52}$$

Rewriting formula (SM.51b) as an SDE provides

$$d\tilde{x} = \left( \overline{xm}N + \frac{\mu}{2} \int_{\mathbb{R}} x\Delta\nu(x,t)dx \right) dt + \|\tilde{x}\|_2 d\tilde{W}_{\tilde{x}}, \tag{SM.53}$$

where

$$d\tilde{W}_{\tilde{x}} = d\hat{\mathbf{W}}_t(\sqrt{Vx^2\nu}) = \frac{1}{\|\tilde{x}\|_2} \int_{\mathbb{R}} x\sqrt{V\nu(x,t)}\dot{W}(x,t)dxdt. \tag{SM.54}$$

Using Itô's quotient rule on $\bar{x} = \tilde{x}/N$, we obtain

$$d\bar{x} = d\left(\frac{\tilde{x}}{N}\right) = \frac{\tilde{x}}{N}\left(\frac{d\tilde{x}}{\tilde{x}} - \frac{dN}{N} - \frac{d\tilde{x}}{\tilde{x}}\frac{dN}{N} + \left(\frac{dN}{N}\right)^2\right) = \frac{d\tilde{x}}{N} - \bar{x}\frac{dN}{N} - \frac{d\tilde{x}}{N}\frac{dN}{N} + \bar{x}\left(\frac{dN}{N}\right)^2. \tag{SM.55}$$

From Table S1 we have $d\tilde{x}dN = \langle \tilde{x}, N \rangle$ and $dN^2 = \|N\|_2^2$. Hence,

$$\begin{aligned} d\bar{x} &= \overline{xm}dt + \frac{\|\tilde{x}\|_2}{N} d\tilde{W}_{\tilde{x}} - \bar{x}\left(\bar{m}dt + \sqrt{\frac{V}{N}}dW_N\right) - \frac{\langle \tilde{x}, N\rangle}{N^2}dt + \bar{x}\frac{\|N\|_2^2}{N^2}dt \\ &= \mathrm{Cov}_t(x,m) + \frac{\|\tilde{x}\|_2}{N}dW_{\tilde{x}} - \bar{x}\sqrt{\frac{V}{N}}dW_N. \end{aligned} \tag{SM.56}$$

Note that

$$\begin{aligned} \frac{\|\tilde{x}\|_2}{N}d\tilde{W}_{\tilde{x}} - \bar{x}\sqrt{\frac{V}{N}}dW_N &= \frac{1}{N}\int_{\mathbb{R}} x\sqrt{V\nu(x,t)}\dot{W}(x,t)dx - \frac{\bar{x}}{N}\int_{\mathbb{R}} \sqrt{V\nu(x,t)}\dot{W}(x,t)dx \\ &= \int_{\mathbb{R}} \frac{x-\bar{x}}{N}\sqrt{V\nu(x,t)}\dot{W}(x,t)dx \end{aligned} \tag{SM.57}$$

and

$$\mathbb{V}\left(\int_{\mathbb{R}} \frac{x-\bar{x}}{N}\sqrt{V\nu(x,t)}\dot{W}(x,t)dx\right) = \frac{V}{N}\int_{\mathbb{R}} (x-\bar{x})^2 p(x,t)dx = V\frac{\sigma^2}{N}. \tag{SM.58}$$

Hence, by setting

$$dW_{\bar{x}} = \frac{\int_{\mathbb{R}} \frac{(x-\bar{x})}{N}\sqrt{V\nu(x,t)}\dot{W}(x,t)dx}{\sqrt{V\sigma^2/N}} \tag{SM.59}$$

we can write

$$d\bar{x} = \mathrm{Cov}_t(x,m)dt + \sqrt{V\frac{\sigma^2}{N}}dW_{\bar{x}}. \tag{SM.60}$$

#### 4.2 Derivation for Trait Variance

We make use of the notation

$$\begin{cases} \|\tilde{\tilde{\sigma}}^2\|_2 = \sqrt{V\int_{\mathbb{R}} x^4\nu(x,t)dx} \\ \langle\tilde{\tilde{\sigma}}^2, N\rangle = V\int_{\mathbb{R}} x^2\nu(x,t)dx = \overline{x^2}VN. \end{cases} \tag{SM.61}$$

Applying formula (SM.51c) provides

$$d\tilde{\tilde{\sigma}}^2 = \left(\overline{x^2m}N + \mu N\right)dt + \|\tilde{\tilde{\sigma}}^2\|_2 d\tilde{\tilde{W}}_{\tilde{\tilde{\sigma}}^2} \tag{SM.62}$$

where

$$d\tilde{\tilde{W}}_{\tilde{\tilde{\sigma}}^2} = d\hat{\mathbf{W}}_t(\sqrt{Vx^4\nu}) = \frac{1}{\|\tilde{\tilde{\sigma}}^2\|_2}\int_{\mathbb{R}} x^2\sqrt{V\nu(x,t)}\dot{W}(x,t)dx. \tag{SM.63}$$

Using Itô's quotient rule on $\overline{x^2} = \tilde{\tilde{\sigma}}^2/N$, we obtain

$$d\overline{x^2} = d\left(\frac{\tilde{\tilde{\sigma}}^2}{N}\right) = \frac{\tilde{\tilde{\sigma}}^2}{N}\left(\frac{d\tilde{\tilde{\sigma}}^2}{\tilde{\tilde{\sigma}}^2} - \frac{dN}{N} - \frac{d\tilde{\tilde{\sigma}}^2}{\tilde{\tilde{\sigma}}^2}\frac{dN}{N} + \left(\frac{dN}{N}\right)^2\right) = \frac{d\tilde{\tilde{\sigma}}^2}{N} - \overline{x^2}\frac{dN}{N} - \frac{d\tilde{\tilde{\sigma}}^2}{N}\frac{dN}{N} + \overline{x^2}\left(\frac{dN}{N}\right)^2. \tag{SM.64}$$

Table S1 implies $d\tilde{\tilde{W}}_{\tilde{\tilde{\sigma}}^2}dW_N = \langle\tilde{\tilde{\sigma}}^2, N\rangle$ and hence

$$\begin{aligned} d\overline{x^2} &= \left(\overline{x^2m} + \mu\right)dt + \frac{\|\tilde{\tilde{\sigma}}^2\|_2}{N}d\tilde{\tilde{W}}_{\tilde{\tilde{\sigma}}^2} - \overline{x^2}\left(\bar{m}dt + \sqrt{\frac{V}{N}}dW_N\right) - \frac{\langle\tilde{\tilde{\sigma}}^2, N\rangle}{N^2}dt + \overline{x^2}\frac{\|N\|_2^2}{N^2}dt \\ &= \left(\overline{x^2m} - \overline{x^2}\bar{m}dt + \mu\right)dt + \frac{\|\tilde{\tilde{\sigma}}^2\|_2}{N}d\tilde{\tilde{W}}_{\tilde{\tilde{\sigma}}^2} - \overline{x^2}\sqrt{\frac{V}{N}}dW_N - \overline{x^2}\frac{V}{N}dt + \overline{x^2}\frac{V}{N}dt \\ &= \left(\mathrm{Cov}_t\left(x^2, m\right) + \mu\right)dt + \frac{\|\tilde{\tilde{\sigma}}^2\|_2}{N}d\tilde{\tilde{W}}_{\tilde{\tilde{\sigma}}^2} - \overline{x^2}\sqrt{\frac{V}{N}}dW_N. \end{aligned} \tag{SM.65}$$

Setting $F(y,z) = y - z^2$, use Itô's formula on $\sigma^2 = F(\overline{x^2}, \bar{x}) = \overline{x^2} - \bar{x}^2$ to obtain:

$$
\begin{aligned}
d\sigma^2 = d\overline{x^2} - 2\bar{x}d\bar{x} - (d\bar{x})^2 = &\left(\mathrm{Cov}_t\left(x^2, m\right) + \mu\right) dt + \frac{\|\tilde{\tilde{\sigma}}^2\|_2}{N} d\tilde{\tilde{W}}_{\tilde{\tilde{\sigma}}^2} - \overline{x^2}\sqrt{\frac{V}{N}} dW_N \\
&- 2\bar{x}\left(\mathrm{Cov}_t(x,m) + \mu dt + \sqrt{\frac{V\sigma^2}{N}} dW_{\bar{x}}\right) - \left(\mathrm{Cov}_t(x,m)dt + \mu dt + \sqrt{\frac{V\sigma^2}{N}} dW_{\bar{x}}\right)^2 \\
= &\left(\mathrm{Cov}_t\left(x^2 - 2\bar{x}x, m\right) + \mu\right) dt + \frac{\|\tilde{\tilde{\sigma}}^2\|_2}{N} d\tilde{\tilde{W}}_{\tilde{\tilde{\sigma}}^2} - \overline{x^2}\sqrt{\frac{V}{N}} dW_N - 2\bar{x}\sqrt{\frac{V\sigma^2}{N}} dW_{\bar{x}} - \left(\frac{V\sigma^2}{N}\right) dt \\
= &\left(\mathrm{Cov}_t\left(x - \bar{x})^2, m\right) + \mu - \frac{V\sigma^2}{N}\right) dt + \frac{\|\tilde{\tilde{\sigma}}^2\|_2}{N} d\tilde{\tilde{W}}_{\tilde{\tilde{\sigma}}^2} - \overline{x^2}\sqrt{\frac{V}{N}} dW_N - 2\bar{x}\sqrt{\frac{V\sigma^2}{N}} dW_{\bar{x}}. \qquad \text{(SM.66)}
\end{aligned}
$$

308 In light of

$$
\begin{aligned}
\frac{\|\tilde{\tilde{\sigma}}^2\|_2}{N} d\tilde{\tilde{W}}_{\tilde{\tilde{\sigma}}^2} - \overline{x^2}\sqrt{\frac{V}{N}} dW_N - 2\bar{x}\sqrt{\frac{V\sigma^2}{N}} dW_{\bar{x}} &= \frac{1}{N}\int_{\mathbb{R}} \left(x^2 - \tilde{\sigma}^2 - 2\bar{x}(x - \bar{x})\right) \sqrt{V\nu(x,t)}\dot{W}(x,t)dx \\
&= \frac{1}{N}\int_{\mathbb{R}} \left((x - \bar{x})^2 - \sigma^2\right) \sqrt{V\nu(x,t)}\dot{W}(x,t)dx \qquad \text{(SM.67)}
\end{aligned}
$$

309 and

$$
\begin{aligned}
\frac{1}{N}\int_{\mathbb{R}} \left(\left((x - \bar{x})^2 - \sigma^2\right) \sqrt{V\nu(x,s)}\right)^2 dx &= \frac{V}{N}\left(\int_{\mathbb{R}} ((x - \bar{x})^4 - 2(x - \bar{x})^2\sigma^2 + \sigma^4)p(x,t)dx\right) \\
&= \frac{V}{N}\left(\overline{(x - \bar{x})^4} - \sigma^4\right) \qquad \text{(SM.68)}
\end{aligned}
$$

310 we set

$$
dW_{\sigma^2} = \frac{\int_{\mathbb{R}} \left((x - \bar{x})^2 - \sigma^2\right) \sqrt{V\nu(x,t)}\dot{W}(x,t)dx}{V\left(\overline{(x - \bar{x})^4} - \sigma^4\right)} \qquad \text{(SM.69)}
$$

311 so that

$$
d\sigma^2 = \mathrm{Cov}_t\left((x - \bar{x})^2, m\right)dt + \left(\mu - V\frac{\sigma^2}{N}\right) dt + \sqrt{V\frac{\overline{(x - \bar{x})^4} - \sigma^4}{N}} dW_{\sigma^2}. \qquad \text{(SM.70)}
$$

312 ## 4.3 Stochastic Dependencies Between $N$, $\bar{x}$ and $\sigma^2$

313 Table S1 implies

$$
dW_N dW_{\bar{x}} = \frac{\int_{\mathbb{R}} (x - \bar{x})\nu(x,t)dx}{\sqrt{N\sigma^2}} dt = 0, \qquad \text{(SM.71a)}
$$

$$dW_N dW_{\sigma^2} = \frac{\int_{\mathbb{R}} \left((x-\bar{x})^2 - \sigma^2\right)\nu(x,t)dx}{\sqrt{\overline{(x-\bar{x})^4} - \sigma^4}} dt = 0, \tag{SM.71b}$$

$$dW_{\bar{x}} dW_{\sigma^2} = \frac{\int_{\mathbb{R}} (x-\bar{x})\left((x-\bar{x})^2 - \sigma^2\right)p(x,t)dx}{\sqrt{\sigma^2\left(\overline{(x-\bar{x})^4} - \sigma^4\right)}} dt = \frac{N\overline{(x-\bar{x})^3}}{\sqrt{\sigma^2\left(\overline{(x-\bar{x})^4} - \sigma^4\right)}} dt. \tag{SM.71c}$$

Hence, stochastic fluctuations in the evolution of abundance $N$ are independent of stochastic fluctuations in the evolution of $\bar{x}$ and $\sigma^2$. However, the stochastic fluctuations in the evolutions of $\bar{x}$ and $\sigma^2$ may be correlated. This is not the case when traits are normally distributed as equation (SM.71c) would then imply $dW_{\bar{x}} dW_{\sigma^2} = 0$.

### 5 Imperfect Inheritance

In this section we provide our approach to model imperfect inheritance, which follows many classical quantitative genetic assumptions.

#### 5.1 Inheritance

To model imperfect heritability we consider the relationship between expressed phenotypes $x \in \mathbb{R}$ and associated genetic values $g \in \mathbb{R}$ known as *breeding values*. The breeding value (called genotypic value in Bulmer, 1971; Walsh and Lynch, 2018) of an individual is the sum of additive effects of the alleles carried by the individual on its expressed trait. Hence, if the trait is encoded by $L$ loci and the additive effect at locus $l$ is $a_l$, then $g = \sum_{l=1}^{L} a_l$. The additive genetic variance $G$ is just the variance of breeding values in a population (Bulmer, 1971; Walsh and Lynch, 2018). Following Lande (1975), we assume a mutation at locus $l$ occurs with probability $M$ and replaces the additive effect $a_l$ with $a_l + \kappa_l$ where $\kappa_l$ is normally distributed with a mean of zero and variance $\mu/M$. Hence, we adopt the Gaussian allelic model of mutation. Next, we implement an infinitesimal approximation by assuming breeding values are determined by an infinite number of loci. Although very general infinitesimal approximations have been provided by Barton et al. (2017), for the sake of simplicity we employ a less technical approach. In particular, we rescale the mutational effects $\kappa_l$ by $1/\sqrt{L}$ and take the limit $L \to \infty$. Then, denoting $g'$ the breeding value of an offspring produced by a parent with breeding value $g$ and $I_l$ the indicator variable determining whether or not a mutation occurs at locus $l$, we have

$$g' = g + \lim_{L\to\infty} \frac{1}{\sqrt{L}} \sum_{l=1}^{L} I_l \kappa_l. \tag{SM.72}$$

This limit implies that $g'$ has expected value $g$ and variance $\mu$. Thus, our assumptions yield the Gaussian descendants approximation coined by Turelli (2017). For a detailed treatment of breeding values, additive genetic variances and more general genetic architectures see Walsh and Lynch (2018).

#### 5.2 Development

Our treatment of the relationship between breeding values and expressed traits follows classical quantitative genetic assumptions such as those used by Bulmer (1971) to investigate the effect of selection on genetic variation. In particular, we ignore epistatic interactions so that effects at different loci combine additively. Since our treatment assumes haploid asexuals, there are no contributions of dominance or inbreeding depression to phenotypic variance. We assume expressed traits for given individuals are normally distributed around their breeding values with a fixed variance $\eta$. Hence, phenotypic variance decomposes as $\sigma^2 = G + \eta$. The variance $\eta$ is referred to as developmental noise (Walsh and Lynch, 2018). For a fixed breeding value $g$, we denote the probability density of a randomly drawn expressed trait $x$ by $\psi(x,g)$ so that

$$\psi(x,g) = \frac{1}{\sqrt{2\pi\eta}} \exp\left(-\frac{(x-g)^2}{2\eta}\right). \tag{SM.73}$$

#### 5.3 Selection On Breeding Values

To include the relationship between breeding values and expressed traits in our framework, we write $\rho(g,t)$ as the abundance density of breeding values at time $t$ so that

$$\int_{-\infty}^{+\infty} \rho(g,t)dg = \int_{-\infty}^{+\infty} \nu(x,t)dx = N(t). \tag{SM.74}$$

We switch our focus from directly modelling the evolution of $\nu(x,t)$ to modelling the evolution of $\rho(g,t)$. Once $\rho(g,t)$ is determined, we can compute $\nu(x,t)$ via

$$\nu(x,t) = \int_{-\infty}^{+\infty} \rho(g,t)\psi(x,g)dg. \tag{SM.75}$$

However, since selection acts on expressed phenotypes, we use the assumed relationship between breeding values and expressed traits to calculate the fitness of breeding values. To motivate the approach taken, consider the problem of inferring the breeding value of an individual given its expressed trait $x$. Denote $\mathfrak{g}$ a random variable representing the unknown breeding value. Under the above model of development we know $x$ is a random sample from a normal distribution with mean $\mathfrak{g}$ and variance $\eta$. Maximizing likelihood suggests $x$ is our best guess for $\mathfrak{g}$, but the actual value of $\mathfrak{g}$ is normally distributed around $x$ with the variance $\eta$. Hence, for fixed $x$, we obtain $\psi(x,g)$ as the probability density of $\mathfrak{g}$. Thus, the mean fitness of a breeding value $g$ across all individuals carrying $g$ can be written as

$$m^*(\rho,g) = \int_{-\infty}^{+\infty} m(\nu,x)\psi(x,g)dx. \tag{SM.76}$$

This is similar to the approach taken by Kimura and Crow (1978) to calculate the overall effects of selection for expressed characters onto the changes in the distribution of alleles encoding those characters. However, instead of focusing on the frequencies of alleles at particular loci, our results focus on the densities of breeding values. With the relationship between $m(\nu,x)$ and $m^*(\rho,g)$ established, we define the evolution of $\rho(g,t)$ by the SPDE

$$\dot{\rho}(g,t) = m^*(\rho,g)\rho(g,t) + \frac{\mu}{2}\frac{\partial^2}{\partial^2 g}\rho(g,t) + \sqrt{V\rho(g,t)}\dot{W}(g,t). \tag{SM.77}$$

Equation (SM.77) is a stochastic generalization of DAGA, the deterministic PDE (4) from §2.1. However, equation (SM.77) describes the evolution of the distribution of breeding values instead of expressed characters. Regardless, whether modelling expressed characters or breeding values, we refer to SPDE of the form (SM.77) as Stochastic Asexual Gaussian allelic models with Abundance dynamics (abbreviated SAGA). In SM §2.1 we develop some heuristics to perform calculations with respect to the space-time white noise term $\dot{W}$ and in SM §3.1 we review the derivation of equation (SM.77) using diffusion-limits of individual-based models.

#### 5.4 Evolution

Assuming $\rho(g,t)$ is Gaussian implies its mode coincides with $\bar{x}$. Furthermore, since $\sigma^2 = G + \eta$, we can use equation (SM.76) and the chain rule from calculus to find

$$\frac{\partial \bar{m}}{\partial G} = \frac{\partial \bar{m}}{\partial \sigma^2}\frac{\partial \sigma^2}{\partial G} = \frac{\partial \bar{m}}{\partial \sigma^2}, \tag{SM.78a}$$

$$\overline{\frac{\partial m}{\partial G}} = \overline{\frac{\partial m}{\partial \sigma^2}\frac{\partial \sigma^2}{\partial G}} = \overline{\frac{\partial m}{\partial \sigma^2}}. \tag{SM.78b}$$

### 6 Derivation of Diffuse Coevolution Model

In this section we provide a derivation of our model of diffuse coevolution driven by resource competition. Since most of the work in this derivation has been completed in SM §4, we focus here on deriving growth rates of each species as a function of trait values and abundance densities of across all species in the community. We then use this fitness function to calculate selection gradients.

#### 6.1 Individual Fitness

We begin with discrete populations of individuals. In particular, we begin by assuming population size $n_i$ is an integer for each species $i = 1, \ldots, S$ before passing to the large population size limit. We assume the competitive effects on fitness for each individual accumulates multiplicatively. For species $i$, the magnitudes of these negative effects increase with the degree of niche-overlap, mediated by the sensitivity $c_i > 0$.

We model niche space using the real line $\mathbb{R}$ and represent locations along this gradient with the symbol $\zeta$. We assume individuals of species $i$ sample the niche gradient following a probability distribution with density $u_i(\zeta, x)$, $x$ being the average niche location sampled or niche center. In particular, we assume individuals sample their environment following a normal distribution so that

$$u_i(\zeta, x) = \frac{U_i}{\sqrt{2\pi w_i}} e^{-\frac{(\zeta - x)^2}{2w_i}}, \tag{SM.79}$$

where $U_i$ represents total niche use (since $U_i = \int u_i(\zeta, x)d\zeta$) and $w_i$ represents niche breadth (the width of the bell curve $u_i$). Following the main text, we define the niche-overlap between individuals of species $i$ and $j$ with trait values $x_i$ and $x_j$ respectively as

$$\mathcal{O}_{ij}(x_i - x_j) = \int_{\mathbb{R}} u_i(\zeta, x_i) u_j(\zeta, x_j) d\zeta = \frac{U_i U_j}{\sqrt{2\pi(w_i + w_j)}} e^{-\frac{(x_i - x_j)^2}{2(w_i + w_j)}}. \tag{SM.80}$$

Denote by $x_{ij}$ the niche center of the $j$-th individual belonging to species $i$. The set of niche centers across all individuals in the community is written $\mathcal{C} = \{x_{ij}\}$. We denote by $\mathfrak{B}_{ij}$ a function that maps $\mathcal{C}$ to the cumulative effect of all competitive interactions on the fitness of the $j$-th individual in species $i$. Since individuals do not compete with themselves the net multiplicative effects on fitness of both interspecific and intraspecific competition on the $j$-th individual in species $i$ can be summarized by

$$\mathfrak{B}_{ij}(\mathcal{C}) = \exp\left(-c_i \sum_{l \neq j} \mathcal{O}_{ii}(x_{ij} - x_{il}) - c_i \sum_{k \neq i} \sum_{l=1}^{n_k} \mathcal{O}_{ik}(x_{ij} - x_{kl})\right). \tag{SM.81}$$

To capture abiotic stabilizing selection we assume resources are normally distributed along the niche gradient. We also assume the concentration of resources is proportional to expected reproductive output. Combining these assumptions, we denote by $e_i(\zeta)$ the fitness benefits for individuals sampling at niche location $\zeta$ so that

$$e_i(\zeta) = Q_i e^{-\frac{A_i}{2}(\theta_i-\zeta)^2}, \tag{SM.82}$$

where $Q_i$ is the maximum expected reproductive output in the absence of competitive interactions, $\theta_i$ is the phenotypic optimum (location along niche axis of most abundant resources) and $A_i > 0$ determines the strength of abiotic stabilizing selection (the sharpness of the resource distribution). Then, we calculate the effect of mismatch between resource use and resource distribution on the fitness of individuals in species $i$ with niche center $x$ as

$$\mathfrak{A}_i(x) = \int_{\mathbb{R}} e_i(\zeta)u_i(\zeta,x)d\zeta = \frac{Q_i U_i}{\sqrt{1+A_i w_i}} e^{-\frac{A_i}{1+A_i w_i}(\theta_i - x)^2}. \tag{SM.83}$$

Writing $\mathfrak{W}_{ij}(\mathcal{C})$ as the average number of offspring left by the $j$-th individual of species $i$, we have

$$\begin{aligned}\mathfrak{W}_{ij}(\mathcal{C}) &= \mathfrak{A}_i(x_{ij})\mathfrak{B}_{ij}(\mathcal{C}) \\ &= \frac{Q_i U_i}{\sqrt{1+A_i w_i}} \exp\left(-\frac{A_i}{1+A_i w_i}(\theta_i - x)^2 - c_i \sum_{l\neq j} \mathcal{O}_{ii}(x_{ij}-x_{il}) - c_i \sum_{k\neq i}\sum_{l=1}^{n_k} \mathcal{O}_{ik}(x_{ij}-x_{kl})\right). \end{aligned} \tag{SM.84}$$

#### 6.2 The Diffusion Limit

To make sense of the diffusion limit, we recall the components of the interacting measure-valued branching process discussed in SM §3.1: (1) the branching rate $\lambda$, (2) the mean $\mathfrak{W}$ and variance $V$ of reproductive output and (3) spatial movement given by Brownian motion with diffusion parameter $\sqrt{\mu}$. For integers $n = 1, 2, \ldots$, we rescale the branching rate by $\lambda \to n$ and fitness by $\mathfrak{W}_{ij}(\mathcal{C}) \to \mathfrak{W}_{ij}^{1/n}(\mathcal{C})$. We rescale the mass of individuals in species $i$ by $N_i(0)/n$ for a fixed positive continuously valued number $N_i(0) > 0$. In the diffusion limit, we take $n \to \infty$. We assume the sequence of initial measures for species $i$, $X_i^{(1)}(0), X_i^{(2)}(0), \ldots$, converges to a limiting measure $\mathfrak{X}_i(0)$ that admits a density $\nu_i(x,0)$ (i.e., $\mathfrak{X}_i(D,0) = \int_D \nu_i(x,0)dx$) such that $\int_{\mathbb{R}}(|x| + x^2 + x^4)\nu_i(x,0)dx < +\infty$ and $\mathfrak{X}_i(\mathbb{R},0) = \int_{\mathbb{R}} \nu_i(x,0)dx = N_i(0) < +\infty$. Hence, rescaled fitness becomes

$$\mathfrak{W}_{ij}^{1/n}(\mathcal{C}) = \mathfrak{A}_i(x_{ij})^{1/n} \exp\left(-\frac{c_i}{n}\frac{N_i(0)}{n}\sum_{l\neq j}\mathcal{O}_{ii}(x_{ij}-x_{il}) - \frac{c_i}{n}\sum_{k\neq i}\frac{N_k(0)}{n}\sum_{l=1}^{n}\mathcal{O}_{1k}(x_{ij}-x_{kl})\right). \tag{SM.85}$$

For large $n$, we have the approximation

$$\mathfrak{W}_{ij}^{1/n}(\mathcal{C}) \approx \mathfrak{A}_i(x_{ij})^{1/n}\left(1 - \frac{c_i}{n}\frac{N_i(0)}{n}\sum_{l\neq j}\mathcal{O}_{ii}(x_{ij}-x_{il}) - \frac{c_i}{n}\sum_{k\neq i}\frac{N_k(0)}{n}\sum_{l=1}^{n}\mathcal{O}_{ik}(x_{ij}-x_{kl})\right). \tag{SM.86}$$

Then, writing $\boldsymbol{\nu} = (\nu_1, \ldots, \nu_S)$, the growth rate of trait value $x_{ij}$ is

$$
\begin{aligned}
m_i(\boldsymbol{\nu}, x_{ij}) &= \lim_{n\to\infty} n\left(\mathfrak{W}_{ij}^{1/n}(\mathcal{C}) - 1\right) \\
&= \lim_{n\to\infty} n\left(\mathfrak{A}_i(x_{ij})^{1/n} - 1\right) - c_i \mathfrak{A}_i(x_{ij})^{1/n}\left(\frac{N_i(0)}{n}\sum_{l\neq j}\mathcal{O}_{ii}(x_{ij}-x_{il}) + \sum_{k\neq i}\frac{N_k(0)}{n}\sum_{l=1}^{n}\mathcal{O}_{ik}(x_{ij}-x_{kl})\right) \\
&= \ln \mathfrak{A}_i(x_{ij}) - c_i\left(\int_{\mathbb{R}} \mathcal{O}_{ii}(x_{ij}-y)\nu_i(y,t)dy + \sum_{k\neq i}\int_{\mathbb{R}} \mathcal{O}_{ik}(x_{ij}-y)\nu_k(y,t)dy\right) \\
&= \ln \mathfrak{A}_i(x_{ij}) - c_i\left(\sum_{k=1}^{S}\int_{\mathbb{R}} \mathcal{O}_{ik}(x_{ij}-y)\nu_k(y,t)dy\right). \qquad \text{(SM.87)}
\end{aligned}
$$

The resulting expression can be used to compute the growth rate for species $i$ associated with any trait value $x \in \mathbb{R}$, which we write as $m_i(\boldsymbol{\nu}, x)$.

#### 6.3 Computing Fitness Gradients

We compute the average niche overlap of an individual in species $i$ with niche location $x$ across all individuals in species $j$ as

$$
\bar{\mathcal{O}}_{ij}(x,t) = \frac{\int_{\mathbb{R}} \mathcal{O}_{ij}(x-y)\nu_j(y,t)dy}{\int_{\mathbb{R}} \nu_j(y,t)dy} = \frac{1}{N_j(t)}\int_{\mathbb{R}} \mathcal{O}_{ij}(x-y)\nu_j(y,t)dy. \qquad \text{(SM.88)}
$$

Following our assumption that individuals of species $i$ sample their environment via a normal distribution with density $u_i(\zeta)$, we further assume normally distributed phenotypes for each of the $S$ species. In this case $\bar{\mathcal{O}}_{ij}(x,t)$ simplifies to

$$
\bar{\mathcal{O}}_{ij}(x,t) = \frac{1}{N_j(t)}\int_{\mathbb{R}} \mathcal{O}_{ij}(x-y)\nu_j(y,t)dy = \frac{U_i U_j}{\sqrt{2\pi(w_i + w_j + \sigma_j{}^2(t))}} \exp\left(-\frac{(x-\bar{x}_j(t))^2}{2(w_i + w_j + \sigma_j{}^2(t))}\right), \qquad \text{(SM.89)}
$$

where $\sigma_i^2(t)$ is the variance of niche-centers in species $i$ at time $t$. Adopting the model of imperfect inheritance formulated in the main text we recall the expressed trait of an individual $x_i$ is normally distributed around its breeding value $g_i$ with variance $\eta_i$. We call $\eta_i$ the variance of environmental deviation and $G_i$, which is the variance of breeding values, the additive genetic variance for species $i$. Under this model of inheritance variance of expressed traits decomposes as $\sigma_i^2(t) = G_i(t) + \eta_i$.

To simplify notation we set

$$
R_i = \ln\left(\frac{Q_i U_i}{\sqrt{1 + A_i w_i}}\right), \qquad \text{(SM.90a)}
$$

$$
a_i = \frac{A_i}{1 + A_i w_i}, \qquad \text{(SM.90b)}
$$

$$
\tilde{b}_{ij}(t) = \frac{1}{w_i + w_j + \sigma_j^2(t)}, \qquad \text{(SM.90c)}
$$

where $R_i$ is the innate growth rate, $a_i$ is the strength of abiotic stabilizing selection and $\tilde{b}_{ij}$ is an intermediate variable mediating the sensitivity of fitness of individuals in species $i$ to interactions with species $j$. With this notation, the growth rate $m_i(\boldsymbol{\nu}, x)$ can be expressed as

$$m_i(\boldsymbol{\nu}, x) = R_i - \frac{a_i}{2}(x-\theta_i)^2 - c_i \sum_{j=1}^{S} N_j U_i U_j \sqrt{\frac{\tilde{b}_{ij}}{2\pi}} \exp\left(-\frac{\tilde{b}_{ij}}{2}(x-\bar{x}_j)^2\right). \tag{SM.91}$$

For the remainder of the derivation we suppress notation indicating dependency on $\boldsymbol{\nu}$ and $x$. From (SM.91) we calculate

$$\frac{\partial m_i}{\partial \bar{x}_i} = c_i N_i U_i^2 \tilde{b}_{ii}(x-\bar{x}_i)\sqrt{\frac{\tilde{b}_{ii}}{2\pi}} \exp\left(-\frac{\tilde{b}_{ii}}{2}(x-\bar{x}_i)^2\right), \tag{SM.92}$$

$$\begin{aligned}\frac{\partial m_i}{\partial G_i} &= \frac{c_i N_i U_i^2}{2}\left(\frac{(x-\bar{x}_i)^2 - G_i - \eta_i - 2w_i}{(G_i+\eta_i+2w_i)^2}\right)\sqrt{\frac{\tilde{b}_{ii}}{2\pi}} \exp\left(-\frac{\tilde{b}_{ii}}{2}(x-\bar{x}_i)^2\right) \\ &= \frac{c_i N_i U_i^2 \tilde{b}_{ii}^2}{2}\left((x-\bar{x}_i)^2 - {\sigma_i}^2 - 2w_i\right)\sqrt{\frac{\tilde{b}_{ii}}{2\pi}} \exp\left(-\frac{\tilde{b}_{ii}}{2}(x-\bar{x}_i)^2\right). \end{aligned} \tag{SM.93}$$

Note that

$$\begin{aligned}&\sqrt{\frac{\tilde{b}_{ii}}{2\pi}} \exp\left(-\frac{\tilde{b}_{ii}}{2}(x-\bar{x}_i)^2\right)\sqrt{\frac{1}{2\pi{\sigma_i}^2}} \exp\left(-\frac{(x-\bar{x}_i)^2}{2{\sigma_i}^2}\right) \\ &\quad = \sqrt{\frac{1}{2\pi({\sigma_i}^2+1/\tilde{b}_{ii})}}\sqrt{\frac{{\sigma_i}^2+1/\tilde{b}_{ii}}{2\pi{\sigma_i}^2/\tilde{b}_{ii}}} \exp\left(-\frac{{\sigma_i}^2+1/\tilde{b}_{ii}}{2{\sigma_i}^2/\tilde{b}_{ii}}(x-\bar{x}_i)^2\right) \\ &\quad = \sqrt{\frac{1}{4\pi({\sigma_i}^2+w_i)}}\sqrt{\frac{2({\sigma_i}^2+w_i)}{2\pi{\sigma_i}^2({\sigma_i}^2+2w_i)}} \exp\left(-\frac{{\sigma_i}^2({\sigma_i}^2+2w_i)}{4({\sigma_i}^2+w_i)}(x-\bar{x}_i)^2\right). \end{aligned} \tag{SM.94}$$

Hence, gradients of the growth rate $m_i$ averaged across the phenotypic distribution $p_i$ become

$$\overline{\frac{\partial m_i}{\partial \bar{x}_i}} = 0, \tag{SM.95}$$

$$\begin{aligned}\overline{\frac{\partial m_i}{\partial G_i}} &= \frac{c_i N_i U_i^2}{2({\sigma_i}^2+2w_i)^2}\left(\frac{({\sigma_i}^2+2w_i){\sigma_i}^2}{2(w_i+{\sigma_i}^2)} - {\sigma_i}^2 - 2w_i\right)\sqrt{\frac{b_{ii}}{2\pi}} \\ &= \frac{c_i N_i U_i^2}{2({\sigma_i}^2+2w_i)}\left(\frac{{\sigma_i}^2}{2({\sigma_i}^2+w_i)} - 1\right)\sqrt{\frac{b_{ii}}{2\pi}} = -\frac{c_i N_i U_i^2 b_{ii}}{2}\sqrt{\frac{b_{ii}}{2\pi}}, \end{aligned} \tag{SM.96}$$

where

$$b_{ij} = \frac{1}{w_i + w_j + {\sigma_i}^2 + {\sigma_j}^2}. \tag{SM.97}$$

The growth rate for species $i$ is

$$\bar{m}_i = R_i - \frac{a_i}{2}\left((\bar{x}_i-\theta_i)^2 + G_i + \eta_i\right) - c_i \sum_{j=1}^{S} N_j U_i U_j \sqrt{\frac{b_{ij}}{2\pi}} \exp\left(-\frac{b_{ij}}{2}(\bar{x}_i-\bar{x}_j)^2\right). \tag{SM.98}$$

Thus, we find the following growth rate gradients

$$\frac{\partial \bar{m}_i}{\partial \bar{x}_i} = a_i(\theta_i - \bar{x}_i) - c_i \sum_{j=1}^{S} N_j U_i U_j b_{ij}(\bar{x}_j - \bar{x}_i)\sqrt{\frac{b_{ij}}{2\pi}} \exp\left(-\frac{b_{ij}}{2}(\bar{x}_i - \bar{x}_j)^2\right), \tag{SM.99}$$

$$\frac{\partial \bar{m}_i}{\partial G_i} = -\frac{a_i}{2} + \frac{c_i}{2}\sum_{j=1}^{S} N_j U_i U_j b_{ij}\left(1 - b_{ij}(\bar{x}_i - \bar{x}_j)^2\right)\sqrt{\frac{b_{ij}}{2\pi}} \exp\left(-\frac{b_{ij}}{2}(\bar{x}_i - \bar{x}_j)^2\right). \tag{SM.100}$$

In particular, we will see

$$\left(\frac{\partial \bar{m}_i}{\partial G_i} - \overline{\frac{\partial m_i}{\partial G_i}}\right) = -\frac{a_i}{2} + \frac{c_i}{2}\left(N_i U_i^2 b_{ii}\sqrt{\frac{b_{ii}}{2\pi}} + \sum_{j=1}^{S} N_j U_i U_j b_{ij}\left(1 - b_{ij}(\bar{x}_i - \bar{x}_j)^2\right)\sqrt{\frac{b_{ij}}{2\pi}} e^{-\frac{b_{ij}}{2}(\bar{x}_i - \bar{x}_j)^2}\right). \tag{SM.101}$$

Applying equations (13a), (17a) and (17b) of the main text recovers system (24) of the main text.

### 7 Competition Coefficients and Selection Gradients

#### 7.1 Definition of Selection Gradients

Our definition of selection gradients differs slightly from traditional definitions. In particular, Lande and Arnold (1983) express the linear selection gradient $\beta$ in general as $\beta = \frac{1}{\sigma^2}\mathrm{Cov}_t(\mathfrak{W}, x)$. This is convenient for discrete time models of mean trait evolution where the change in mean trait between generations is captured by

$$\Delta \bar{x} = \frac{G}{\sigma^2}\mathrm{Cov}_t(\mathfrak{W}, x) = G\beta. \tag{SM.102}$$

However, in our case, we model mean trait evolution in continuous time via

$$\frac{d\bar{x}}{dt} = \frac{G}{\sigma^2}\mathrm{Cov}_t(m, x). \tag{SM.103}$$

Hence, we define the linear selection gradient $\beta := \frac{1}{\sigma^2}\mathrm{Cov}_t(m, x)$. Similarly, the quadratic selection gradient $\gamma$ is expressed in Lande and Arnold (1983) as $\gamma = \frac{1}{\sigma^4}\mathrm{Cov}_t(\mathfrak{W}, (x - \bar{x})^2)$. Then, in analogy to our definition of $\beta$, we define $\gamma := \frac{1}{\sigma^4}\mathrm{Cov}_t(m, (x - \bar{x})^2)$.

#### 7.2 Selection Gradients Under Abiotic Stabilizing Selection and Resource Competition

Here we derive expressions for selection gradients under our model of diffuse coevolution driven by resource competition. Combining our definitions of selection gradients with the results found in SM §6, our model of diffuse coevolution yields, for species $i$,

$$\beta_i = a_i(\theta_i - \bar{x}_i) - c_i \sum_{j=1}^{S} N_j U_i U_j b_{ij}(\bar{x}_j - \bar{x}_i)\sqrt{\frac{b_{ij}}{2\pi}} e^{-\frac{b_{ij}}{2}(\bar{x}_j - \bar{x}_i)^2}, \tag{SM.104a}$$

$$\gamma_i = -a_i + c_i \left( N_i U_i^2 b_{ii} \sqrt{\frac{b_{ii}}{2\pi}} + \sum_{j=1}^{S} N_j U_i U_j b_{ij} \left(1 - b_{ij}(\bar{x}_i - \bar{x}_j)^2\right) \sqrt{\frac{b_{ij}}{2\pi}} e^{-\frac{b_{ij}}{2}(\bar{x}_i - \bar{x}_j)^2} \right). \tag{SM.104b}$$

Note these selection gradients can be additively partitioned as $\beta_i = \beta_i^{(a)} + \sum_{j=1}^{S} \beta_{ij}$ and $\gamma_i = \gamma_i^{(a)} + \sum_{j=1}^{S} \gamma_{ij}$ where $\beta_i^{(a)}, \gamma_i^{(a)}$ denote the components due to abiotic stabilizing selection and $\beta_{ij}, \gamma_{ij}$ denote the components due to interactions with species $j$. In particular, we find $\beta_i^{(a)} = a_i(\theta_i - \bar{x}_i)$, $\gamma_i^{(a)} = -a_i$ and

$$\beta_{ij} = c_i N_j U_i U_j b_{ij} (\bar{x}_i - \bar{x}_j) \sqrt{\frac{b_{ij}}{2\pi}} e^{-\frac{b_{ij}}{2}(\bar{x}_i - \bar{x}_j)^2}, \tag{SM.105a}$$

$$\gamma_{ij} = c_i N_j U_i U_j b_{ij} \left(1 - b_{ij}(\bar{x}_i - \bar{x}_j)^2\right) \sqrt{\frac{b_{ij}}{2\pi}} e^{-\frac{b_{ij}}{2}(\bar{x}_i - \bar{x}_j)^2}, \ i \neq j \tag{SM.105b}$$

$$\gamma_{ii} = 2 c_i N_i U_i^2 b_{ii} \sqrt{\frac{b_{ii}}{2\pi}}, \ i = j. \tag{SM.105c}$$

#### 7.3 High Richness Approximations for Moments of Competition Coefficients and Selection Gradients

Here we derive covariances between competition coefficients and selection gradients following the model of diffuse coevolution derived in SM §6. We assume the community is very rich (i.e., the number of species $S$ is very large) and that the distribution of mean traits is approximately normal and independent of the distribution of abundance. We denote by $\bar{\bar{x}}$, $V_{\bar{x}}$ the community-wide mean and variance of species mean traits and by $\bar{N}, V_N$ the community-wide mean and variance of species abundances. For simplicity we assume constant species trait variances and niche breadths so that $\sigma_i^2 = \sigma^2$ and $w_i = w$ for some $\sigma^2, w > 0$. Thus $b_{ij} = b = 1/(2\sigma^2 + 2w)$ for each $i, j = 1, \ldots, S$. Under these conditions, we can express competition coefficients, linear selection gradients and quadratic selection gradients respectively as

$$\alpha_{ij} = c_i U_i U_j \sqrt{\frac{b}{2\pi}} e^{-\frac{b}{2}(\bar{x}_i - \bar{x}_j)^2}, \tag{SM.106a}$$

$$\beta_{ij} = c_i U_i U_j N_i b (\bar{x}_i - \bar{x}_j) \sqrt{\frac{b}{2\pi}} e^{-\frac{b}{2}(\bar{x}_i - \bar{x}_j)^2}, \tag{SM.106b}$$

$$\gamma_{ij} = c_i U_i U_j N_i b \left(1 - b(\bar{x}_i - \bar{x}_j)^2\right) \sqrt{\frac{b}{2\pi}} e^{-\frac{b}{2}(\bar{x}_i - \bar{x}_j)^2}, \ i \neq j. \tag{SM.106c}$$

To compute statistical distributions of these quantities we draw $i$ and $j$ independently from the set $\{1, \ldots, S\}$ each with probability $1/S$. Then the event $i = j$ occurs with probability $1/S^2$. We suppose $S$ is large enough that we can safely ignore the event $i = j$.

Under our model of diffuse coevolution, the competition coefficients and selection gradients can be written in terms of the difference $D_{ij} = \bar{x}_i - \bar{x}_j$. By our assumption that $i$ and $j$ are drawn independently and that $\bar{x}_i$, $\bar{x}_j$ approximately follow a normal distribution with mean $\bar{\bar{x}}$ and variance $V_{\bar{x}}$, we see the distribution of $D_{ij}$ is approximated by a normal distribution with mean zero and variance $2V_{\bar{x}}$.

We suppose the strengths of competition $c_i$ and niche-use parameters $U_i$ are distributed independently of mean traits, abundances and each other. We write $\bar{c}$, $\bar{U}$ and $V_c$, $V_U$ as the mean and variance of these parameters respectively.

##### 7.3.1 Means and Variances of Competition Coefficients and Selection Gradients

Combining the above assumptions and notation, we can approximate the expectations of competition coefficients and selection gradients via

$$\bar{\alpha} = \frac{1}{S^2}\sum_{i,j=1}^{S} \alpha_{ij} \approx \bar{c}\bar{U}^2 \int_{\mathbb{R}} \sqrt{\frac{b}{2\pi}} e^{-\frac{b}{2}D^2} \frac{1}{\sqrt{4\pi V_{\bar{x}}}} e^{-\frac{D^2}{4V_{\bar{x}}}} dD = \bar{c}\bar{U}^2 \sqrt{\frac{b}{2\pi(2V_{\bar{x}}b+1)}}, \tag{SM.107a}$$

$$\bar{\beta} = \frac{1}{S^2}\sum_{i,j=1}^{S} \beta_{ij} \approx \bar{c}\bar{U}^2\bar{N}b \int_{\mathbb{R}} D\sqrt{\frac{b}{2\pi}} e^{-\frac{b}{2}D^2} \frac{1}{\sqrt{4\pi V_{\bar{x}}}} e^{-\frac{D^2}{4V_{\bar{x}}}} dD = 0, \tag{SM.107b}$$

$$\begin{aligned}\bar{\gamma} = \frac{1}{S^2}\sum_{i,j=1}^{S} \gamma_{ij} &\approx \bar{c}\bar{U}^2\bar{N}b \int_{\mathbb{R}} \left(1 - bD^2\right) \sqrt{\frac{b}{2\pi}} e^{-\frac{b}{2}D^2} \frac{1}{\sqrt{4\pi V_{\bar{x}}}} e^{-\frac{D^2}{4V_{\bar{x}}}} dD \\ &= \bar{c}\bar{U}^2\bar{N}b \sqrt{\frac{b}{2\pi(2V_{\bar{x}}b+1)}} \left(\frac{1}{2V_{\bar{x}}b+1}\right) = \frac{\bar{\alpha}\bar{N}b}{2V_{\bar{x}}b+1}. \end{aligned} \tag{SM.107c}$$

Similarly, their variances can be approximated as

$$\begin{aligned}\mathrm{Var}(\alpha) = \overline{\alpha^2} - \bar{\alpha}^2 &\approx (V_c + \bar{c}^2)(V_U + \bar{U}^2)^2 \int_{\mathbb{R}} \frac{b}{2\pi} e^{-bD^2} \frac{1}{\sqrt{4\pi V_{\bar{x}}}} e^{-\frac{D^2}{4V_{\bar{x}}}} dD - \bar{\alpha}^2 \\ &= \frac{(V_c + \bar{c}^2)(V_U + \bar{U}^2)^2 b}{2\pi\sqrt{4V_{\bar{x}}b+1}} - \bar{\alpha}^2, \end{aligned} \tag{SM.108a}$$

$$\begin{aligned}\mathrm{Var}(\beta) = \overline{\beta^2} - \bar{\beta}^2 &\approx (V_c + \bar{c}^2)(V_U + \bar{U}^2)^2(V_N + \bar{N}^2)b \int_{\mathbb{R}} D^2 \frac{b}{2\pi} e^{-bD^2} \frac{1}{\sqrt{4\pi V_{\bar{x}}}} e^{-\frac{D^2}{4V_{\bar{x}}}} dD \\ &= \frac{(V_c + \bar{c}^2)(V_U + \bar{U}^2)^2(V_N + \bar{N}^2)b^2 V_{\bar{x}}}{\pi(4V_{\bar{x}}b+1)^{3/2}}, \end{aligned} \tag{SM.108b}$$

$$\begin{aligned}\mathrm{Var}(\gamma) = \overline{\gamma^2} - \bar{\gamma}^2 &\approx (V_c + \bar{c}^2)(V_U + \bar{U}^2)^2(V_N + \bar{N}^2)b \int_{\mathbb{R}} \left(1 - bD^2\right)^2 \frac{b}{2\pi} e^{-bD^2} \frac{1}{\sqrt{4\pi V_{\bar{x}}}} e^{-\frac{D^2}{4V_{\bar{x}}}} dD - \bar{\gamma}^2 \\ &= \frac{(V_c + \bar{c}^2)(V_U + \bar{U}^2)^2(V_N + \bar{N}^2)b^2}{\pi\sqrt{4V_{\bar{x}}b+1}} \left(1 - 2b\left(\frac{V_{\bar{x}}}{4V_{\bar{x}}b+1}\right) + 3b^2\left(\frac{V_{\bar{x}}}{4V_{\bar{x}}b+1}\right)^2\right) - \bar{\gamma}^2. \end{aligned} \tag{SM.108c}$$

##### 7.3.2 Mean and Variance of Absolute Values of Linear Selection Gradients

Since our above assumptions imply a certain degree of symmetry across the community, we find the average linear selection gradient is zero. Then, to extract information about the total quantity of linear selection occurring in the community, we consider the absolute values of linear selection gradients. Following the above assumptions we can express $|\beta_{ij}|$ as

$$|\beta_{ij}| = c_i U_i U_j N_i b |D_{ij}| \sqrt{\frac{b}{2\pi}} e^{-\frac{b}{2}D_{ij}^2}. \tag{SM.109}$$

The mean of $|\beta_{ij}|$ can then be approximated as

$$\overline{|\beta|} \approx \bar{c}\bar{U}^2\bar{N}b\int_{\mathbb{R}}|D|\sqrt{\frac{b}{2\pi}}e^{-\frac{b}{2}D^2}\frac{1}{\sqrt{4\pi V_{\bar{x}}}}e^{-\frac{D^2}{4V_{\bar{x}}}}dD$$
$$= \bar{c}\bar{U}^2\bar{N}b\sqrt{\frac{b}{2\pi(2V_{\bar{x}}b+1)}}\int_{\mathbb{R}}|D|\sqrt{\frac{b}{2\pi(2V_{\bar{x}}b+1)}}e^{-\frac{2V_{\bar{x}}b+1}{2b}D^2}dD. \quad \text{(SM.110)}$$

Computing the integral on the RHS is equivalent to computing the mean of the absolute value of a normally distributed random variable with mean zero and variance $\frac{b}{2V_{\bar{x}}b+1}$. It is well known that the absolute value $|Z|$ of a normally distributed random variable $Z$, itself taking mean zero and variance $V_Z$, has mean $\overline{|Z|} = \sqrt{2V_Z/\pi}$. Hence, we can use this information to compute

$$\overline{|\beta|} \approx \bar{c}\bar{U}^2\bar{N}b\sqrt{\frac{b}{2\pi(2V_{\bar{x}}b+1)}}\sqrt{\frac{2b}{\pi(V_{\bar{x}}b+1)}} = \frac{\bar{c}\bar{U}^2\bar{N}b^2}{\pi(2V_{\bar{x}}b+1)}. \quad \text{(SM.111)}$$

The variance $\text{Var}(|\beta|)$ is a bit easier to calculate. In particular, we can approximate the variance of absolute values of $\beta$ via

$$\text{Var}(|\beta|) = \overline{|\beta|^2} - \overline{|\beta|}^2 = \overline{\beta^2} - \overline{|\beta|}^2 \approx \text{Var}(\beta) - \overline{|\beta|}^2, \quad \text{(SM.112)}$$

where we have capitalized on the result $\bar{\beta} \approx 0$.

##### 7.3.3 Correlations Between Competition Coefficients and Selection Gradients

Following the above assumptions and notation, the covariance of competition coefficients $\alpha_{ij}$ and linear selection gradients $\beta_{ij}$ can be approximated as

$$\text{Cov}(\alpha,\beta) = \overline{\alpha\beta} - \bar{\alpha}\bar{\beta} \approx (V_c+\bar{c}^2)(V_U+\bar{U}^2)^2\bar{N}b\int_{\mathbb{R}}D\frac{b}{2\pi}e^{-bD^2}\frac{1}{\sqrt{4\pi V_{\bar{x}}}}e^{-\frac{D^2}{4V_{\bar{x}}}}dD = 0. \quad \text{(SM.113)}$$

Again, this result follows from our assumptions on the distribution of model parameters across the community. To extract information about the covariance between competition coefficients and the magnitude of linear selection, we compute $\text{Cov}(\alpha, |\beta|)$. This quantity can be approximated by

$$\text{Cov}(\alpha,|\beta|) = \overline{\alpha|\beta|} - \bar{\alpha}\overline{|\beta|} \approx (V_c+\bar{c}^2)(V_U+\bar{U}^2)^2\bar{N}b\int_{\mathbb{R}}|D|\frac{b}{2\pi}e^{-bD^2}\frac{1}{\sqrt{4\pi V_{\bar{x}}}}e^{-\frac{D^2}{4V_{\bar{x}}}}dD - \bar{\alpha}\overline{|\beta|}$$
$$= \frac{(V_c+\bar{c}^2)(V_U+\bar{U}^2)^2\bar{N}b^2}{\pi(4V_{\bar{x}}b+1)}\sqrt{\frac{V_{\bar{x}}}{\pi}} - \bar{\alpha}\overline{|\beta|}, \quad \text{(SM.114)}$$

where we have again made use of the properties of absolute values of normally distributed random variables.

The covariance between competition coefficients and quadratic selection gradients can be approximated by

$$\text{Cov}(\alpha,\gamma) = \overline{\alpha\gamma} - \bar{\alpha}\bar{\gamma} \approx (V_c+\bar{c}^2)(V_U+\bar{U}^2)^2\bar{N}b\int_{\mathbb{R}}(1-bD^2)\frac{b}{2\pi}e^{-bD^2}\frac{1}{\sqrt{4\pi V_{\bar{x}}}}e^{-\frac{D^2}{4V_{\bar{x}}}}dD - \bar{\alpha}\bar{\gamma}$$
$$= \frac{(V_c+\bar{c}^2)(V_U+\bar{U}^2)^2\bar{N}b^2(2V_{\bar{x}}b+1)}{2\pi(4V_{\bar{x}}b+1)^{3/2}} - \bar{\alpha}\bar{\gamma}. \quad \text{(SM.115)}$$

Finally, these approximations can be used to approximate the correlations between competition coefficients and selection gradients via

$$\mathrm{Corr}(\alpha, |\beta|) = \frac{\mathrm{Cov}(\alpha, |\beta|)}{\sqrt{\mathrm{Var}(\alpha)\mathrm{Var}(|\beta|)}}, \tag{SM.116a}$$

$$\mathrm{Corr}(\alpha, \gamma) = \frac{\mathrm{Cov}(\alpha, \gamma)}{\sqrt{\mathrm{Var}(\alpha)\mathrm{Var}(\gamma)}}. \tag{SM.116b}$$
